# Expanded gene targeting in RNA hacking with G-tract-supply Staple oligomer

**DOI:** 10.1101/2025.06.17.660051

**Authors:** Tomoki Kida, Yua Hasegawa, Miko Kato, Mahiro Ohtani, Ayu Matsumoto, Ririka Taniguchi, Rinka Ohno, Hiroshi Sugiyama, Toshihiro Ihara, Masaki Hagihara, Shin-ichi Sato, Yousuke Katsuda

## Abstract

RNA hacking (RNAh) is a gene regulation technology that employs a short oligonucleotide, termed a Staple oligomer, to induce the formation of RNA G-quadruplex structures on target mRNAs. While RNAh has the potential to target approximately 65% of human mRNAs, its applicability to the remaining genes is restricted by the sequence constraints. Herein, we present the G-tract-supply Staple oligomer (Gs-Staple oligomer), designed to expand the range of targetable mRNAs within the RNAh framework. Incorporating G-tracts into Staple oligomers alleviates the sequence constraints, enabling access to a broader range of mRNA targets. Gs-Staple oligomers effectively suppressed the translation of target proteins in mammalian cells and in vivo. Furthermore, the gene suppression could be precisely modulated by adjusting the linker length between the G-tracts. These findings have significantly expanded the versatility of RNAh, suggesting its potential for further development while highlighting its potential to be utilized as a nucleic acid-based tool for research and clinical medicine.

---

Unlike conventional protein-targeted drugs, nucleic acid-based therapeutics directly target DNA or RNA and thus have superior target specificity^1-5^. The design of seed molecules of nucleic acid-based therapeutics is a relatively straightforward and rational process that relies on the nucleotide sequence of the target gene. This makes nucleic acid-based therapeutics a promising approach for treating rare and severe diseases. Small interfering RNA (siRNA), which induce RNA interference (RNAi), and antisense oligonucleotides (ASOs), which trigger RNaseH-mediated degradation of target mRNA, represent efficacious nucleic acid-based tools for gene silencing^6-8^. These technologies have the potential to be highly effective in treating a wide range of diseases, including genetic disorders and cancers^9^. However, there are a number of hurdles that must be overcome before these technologies can be utilized in the field of nucleic acid therapeutics. These issues include the introduction of chemical modifications and the minimization of off-target effects^10-15^. For example, to enhance the stability of these oligonucleotides for *in vivo* medical applications, various chemical modifications and non-natural nucleic acids have been introduced into the nucleic acid components^16, 17^. While these chemical modifications enhance *in vivo* stability, they also have the potential to disrupt essential biological processes mediated by Argonaute or RNaseH, leading to a substantial loss of gene silencing efficiency^18-21^. Another significant concern is the prevalence of off-target effects on non-target mRNAs which restrict the practical application of siRNA and ASO technologies to a limited number of cases.

We have recently reported RNA hacking (RNAh) as a technology which induces RNA G-quadruplex (rG4) formation on target mRNA by use of a Staple oligomer (Fig. 1a)^22^. The Staple oligomer binds to two specific locations in the proximity of four G-tracts on target mRNA. Consequently, the G-tracts are brought into proximity, and the formation of a rG4 is induced. rG4 is a thermodynamically stable structure that, when positioned upstream on mRNA, can impede ribosome progression and thereby suppress protein translation^23^. The gene suppression mechanism by which Staple oligomers function is entirely distinct from that of siRNA and ASOs in that the operation of Staple oligomers is independent of the activity of endogenous enzymes. This unique mechanism enables gene suppression levels to be maintained even when non-natural nucleic acids are incorporated as components of the Staple oligomer. In addition, a notable advantage of RNAh is its capacity to minimize off-target effects. In contrast to the mechanism of siRNAs and ASOs, which bind to a single site on the target mRNA and subsequently activate the gene-silencing process with relevant enzymes, the mechanism of Staple oligomers requires two distinct activation conditions. First, Staple oligomers must bind simultaneously to two separate and specific sequences on mRNA. Second, these sequences must be near four G-tracts that are brought into close proximity by binding the Staple to form the G-quadruplex (G4) structure. Consequently, the off-target effects are significantly reduced compared to those observed with siRNAs or ASOs. Bioinformatics research estimates predict that RNAh technology using Staple oligomers has the potential to target approximately 65% of known human mRNAs. Sequence constraints arise when there are an insufficient number of G-tracts in the target mRNA. To enhance the versatility of RNAh, it is imperative for Staple oligomers to be able to compensate for the lack of G-tracts to trigger G4 formation.

**Fig. 1.**
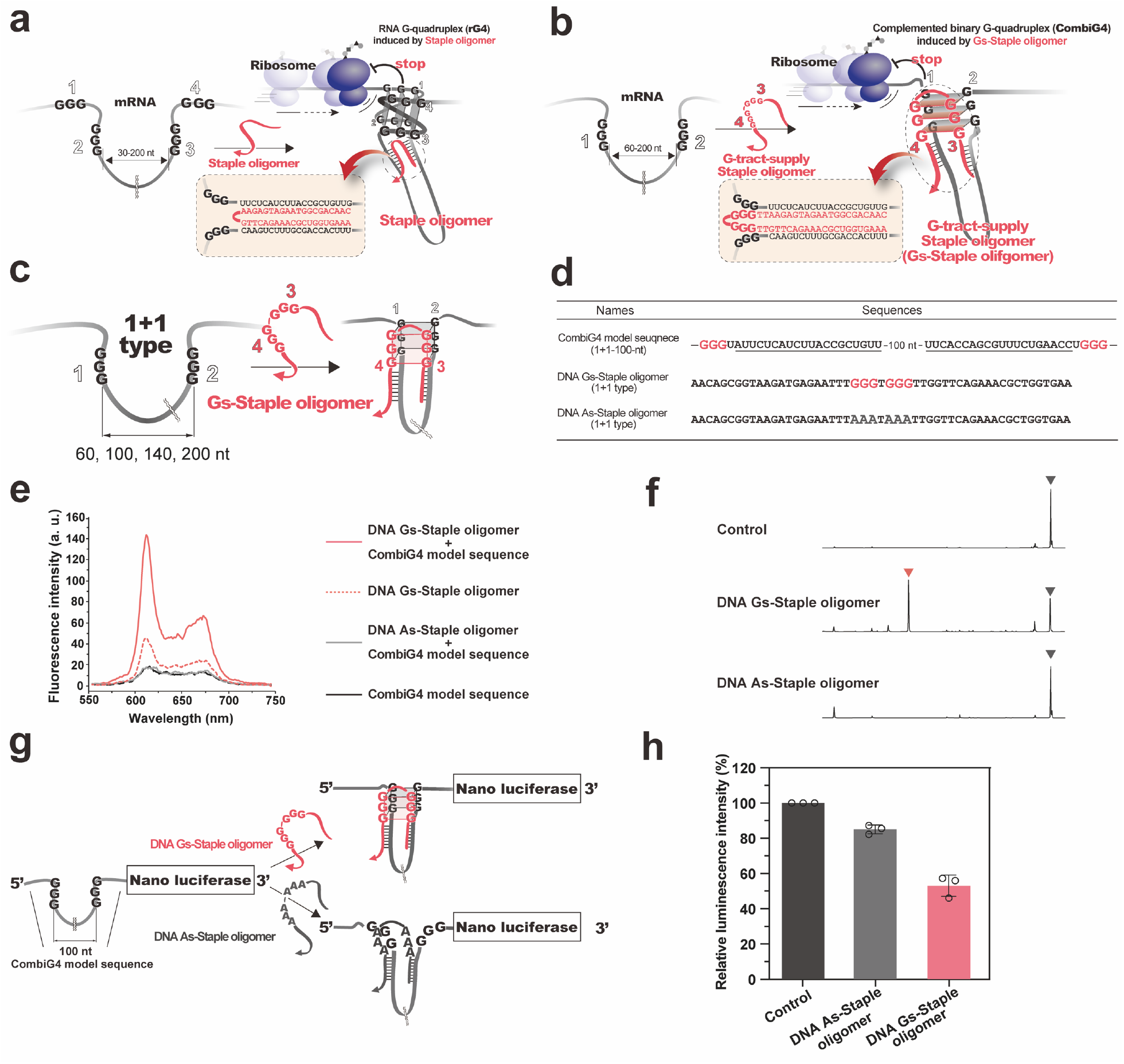
Characterizing G-tracts-supply Staple oligomer. **a,** Mechanism of action of predecessor Staple oligomers which induce rG4 formation. **b,** Mechanism of action of G-tract-supply Staple oligomers (Gs-Staple oligomers) which induce combiG4 formation. **c,** 1+1 type CombiG4 model sequences are illustrated. CombiG4 model sequences have two G-tracts that are split into a 1+1 arrangement with 60, 100, 140, or 200-nt loops. **d,** The sequences of 1+1_100-nt combiG4 model, DNA Gs-Staple oligomer, and DNA As-Staple oligomer. The G-tracts are shown in red. The solid underlines mark the DNA Gs-Staple oligomer or DNA As-Staple oligomer recognition sites. **e,** Evaluation of DNA/RNA combiG4 formation by Gs-Staple oligomer with NMM. The red, red-dashed, gray, and black curves show the fluorescence emission spectra of NMM in the presence of the DNA Gs-Staple oligomer with the combiG4 model sequence RNA, the Gs-Staple oligomer alone, the As-Staple oligomer with the combiG4 model sequence, or the combiG4 model sequence RNA alone, respectively. **f,** RTase stop assay spectra of the 1+1_100-nt combiG4 model sequence. RTase-mediated cDNA synthesis was interrupted on combiG4 model sequences in the presence of each DNA Gs-Staple oligomer. The red arrowhead marks the arrest site of RTase with DNA/RNA combiG4 formation induced by the DNA Gs-Staple oligomer, and the black arrowheads mark the end of RTase elongation without induction of DNA/RNA combiG4 formation. **g,** The 1+1_100-nt combiG4 model sequence was introduced to the 5’ UTR of Nanoluc gene. The protein expression level was determined by Nanoluc luminescence intensity. DNA Gs-Staple oligomer induces DNA/RNA combiG4 formation (upper image), whereas DNA As-Staple oligomer does not (lower image). **h,** Effect of the DNA Gs-Staple oligomer or the As-Staple oligomer on the translation of reporter RNA *in vitro*. Data are the mean ± S.D. of three independent experiments.

It was recently reported that incorporating up to three G-tracts into ASOs can compensate for the lack of G-tracts in the target mRNA, thereby promoting the formation of a stable heteroduplex through a hybrid rG4 structure. This approach is referred to as G-tract-supply antisense (g-AS)^24-27^. Despite the notable efficacy of the g-AS strategy, its simple working principle–based on complementarity to a single site of the target sequence–raises concerns about potential off-target effects akin to those observed with the use of siRNA and ASOs. Here, we describe the G-tract-supply Staple oligomer (Gs-Staple oligomer), which incorporates G-tracts within its sequence to form the G4 structure. We use the term complemented binary G4 (combiG4) structure to refer to a G4 structure formed between a target mRNA and an exogenous oligonucleotide (DNA or RNA Gs-Staple oligomer) which supplements the G-tracts necessary for G4 formation. The integration of G-tract sequences within the Gs-Staple oligomer serves to alleviate the sequence constraints that currently limit the applicability of RNAh technology (Fig. 1b). Gs-Staple oligomers work in a manner analogous to existing Staple oligomers, thereby minimizing off-target effects. Gs-Staple oligomers offer versatility comparable to that of existing tools such as siRNA and ASOs, with the additional benefits of unrestricted incorporation of non-natural nucleic acids and the minimization of off-target effects. Accordingly, it presents a promising advance in the field of nucleic acid-based therapeutics.

## Results

### Induction of combiG4 formation by Gs-Staple oligomer

Gs-Staple oligomers have been engineered to hybridize to two distinct regions on a target mRNA, forming a combiG4 with G-tracts on RNA. We first constructed a series of model sequences as mRNA targets to investigate this mechanism. These sequences containing two G-tracts separated by 60, 100, 140, or 200 nucleotides were designated as model sequences that form 1+1 type combiG4 structures with the target sequence (Fig. 1c-d and Table S1). The DNA Gs-Staple oligomer – comprised of two G-tracts separated by a one-nucleotide thymidine linker – was prepared to target the model sequences, along with the As-Staple oligomer in which the G-tracts were replaced with A-tracts as a negative control (Fig. 1d and S1a). To confirm the formation of DNA/RNA combiG4 structures by coordination with the G-tracts contained in the DNA Gs-Staple oligomer and those of the target RNA, we performed fluorescence measurements using N-methylmesoporphyrin IX (NMM), which exhibits enhanced fluorescence at 610 nm upon binding G4 structures^28-30^. A significant increase in fluorescence was observed for all model sequences upon the addition of the DNA Gs-Staple oligomer. In contrast, fluorescence enhancement was detected neither when the DNA As-Staple oligomer was added to the model sequences nor when the Gs-Staple DNA oligomer was evaluated on its own (Fig. 1e and S1b). These observations provide evidence that the G-tracts in the DNA Gs-Staple oligomer and those in the model sequence coalesce to form a DNA/RNA combiG4 structure.

It has been demonstrated that G4 structures formed on target RNA impede the elongation of cDNA in reverse transcription. Hagihara et al. developed the RTase stop assay for the detection and identification of G4 structures^24, 31^. The RTase stop assay was employed to assess the DNA/RNA combiG4 formation induced by the DNA Gs-Staple oligomer. The introduction of the DNA Gs-Staple oligomer impeded the progression of reverse transcription in all 1+1 type combiG4 model sequences (Fig. 1f and S2a). In contrast, the DNA As-Staple oligomer did not affect the formation of full-length cDNA during reverse transcription to (Fig. 1f and S2a). To confirm that reverse transcription was impeded by the DNA/RNA combiG4 structure, the RTase stop assay was conducted in a buffer solution containing lithium ions instead of potassium ions. The thermodynamic stability of G4 structures depends on the size of the cations: potassium ions provide high stability, whereas the presence of lithium ions results in low stability^32^. As anticipated, the introduction of the DNA Gs-Staple oligomer did not impede the progression of reverse transcription in the presence of lithium ions (Fig. S2b). Subsequently, an *in vitro* translation assay was conducted to assess the potential inhibitory activity of the DNA Gs-Staple oligomer on protein translation. The suppression efficiency of the DNA Gs-Staple oligomer on translation was evaluated using a Nano luciferase (NanoLuc) reporter gene containing the 1+1_100-nt combiG4 model sequence, which consists of two G-tracts separated by a 100-nt intervening sequence in its 5’ untranslated region (UTR) (Fig. 1g and Table S1). Introduction of the Gs-Staple oligomer resulted in a reduction of NanoLuc translation. In contrast, addition of the DNA As-Staple oligomer exhibited no detectable decrease in Nanoluc expression (Fig. 1h). These findings suggest that the DNA/RNA combiG4 formed by the G-tracts on the Gs-Staple oligomer and those on the target RNA can suppress protein translation.

In case the target mRNA has less than four G-tracts available for G4 structure formation, it is necessary to use Gs-Staple oligomers with a complementary number of G-tracts. As a new variety, we experimented with a model target with three G-tracts divided into 2+1 with 60-, 100-, 140-, and 200-nt intervening sequences and the DNA Gs-Staple with one G-tract (Fig. S3a-b, Table S1). These constructs were evaluated with the corresponding Gs-Staple and As-Staple oligomers using experimental techniques parallel to those previously described for the 1+1 model target. NMM fluorescence measurements revealed a significant signal increase only in the presence of both the Gs-Staple oligomer and the 2+1 type combiG4 model sequence (Fig. S3c). RTase stop assay results confirmed that reverse transcription was arrested at the combiG4 site in the presence of the Gs-Staple oligomer. In contrast, the inhibition of reverse transcription was not observed in the presence of the As-Staple oligomer (Fig. S3d). RTase stop assay in a lithium-containing buffer showed that the DNA Gs-Staple oligomer did not impede the progression of reverse transcription (Fig. S3e). Furthermore, *in vitro* translation was performed using a NanoLuc reporter containing a 2+1_100-nt combiG4 model sequence in the 5’ UTR (Fig. S3f and Table S1). The Gs-Staple oligomer once again diminished NanoLuc translation, while the DNA As-Staple oligomer exerted no discernible influence (Fig. S4g). These results show that Gs-Staple oligomers can be logically designed according to the number of G-tracts present on the target mRNA, suggesting that the RNAh technology can be used to target a wide range of mRNAs by tailoring appropriate G4 structures.

### Exploration of linker length effects between G-tracts in Gs-Staple oligomers

We modified the number of linker bases between the G-tracts to optimize the DNA Gs-Staple oligomer with two G-tracts. To this end, we prepared four Gs-Staple oligomers with linkers containing 1, 2, 4, and 8 dT_n_ units between the G-tracts, targeting 1+1 type RNA model sequences with 60-, 100-, 140-, or 200-nt intervening loop regions (Fig. 2a, Table S1, and S2). First, the effect of linker length on the formation of DNA/RNA combiG4 structures was assessed using NMM. The Gs-Staple oligomer with the 1-nt linker exhibited the highest fluorescence intensity, which decreased progressively with increasing linker length (Fig. 2b and S4a). Subsequently, melting experiments were conducted to evaluate the impact of linker length on the thermal stability of the combiG4 structures by monitoring the fluorescence change of a FAM label attached to the 5’-end of the DNA Gs-Staple oligomer. As anticipated, the DNA Gs-Staple oligomer with the 1-nt linker showed the highest melting temperature, and the stability decreased as the linker length increased (Fig. S4b). The thermal stability of the Gs-Staple oligomer with a linker of 4-nt or 8-nt was found to be similar to that of the DNA As-Staple oligomer. These findings suggest that the linker length between the G-tracts in the Gs-Staple oligomer affects the thermal stability of the combiG4 complexes with target RNA.

**Fig. 2.**
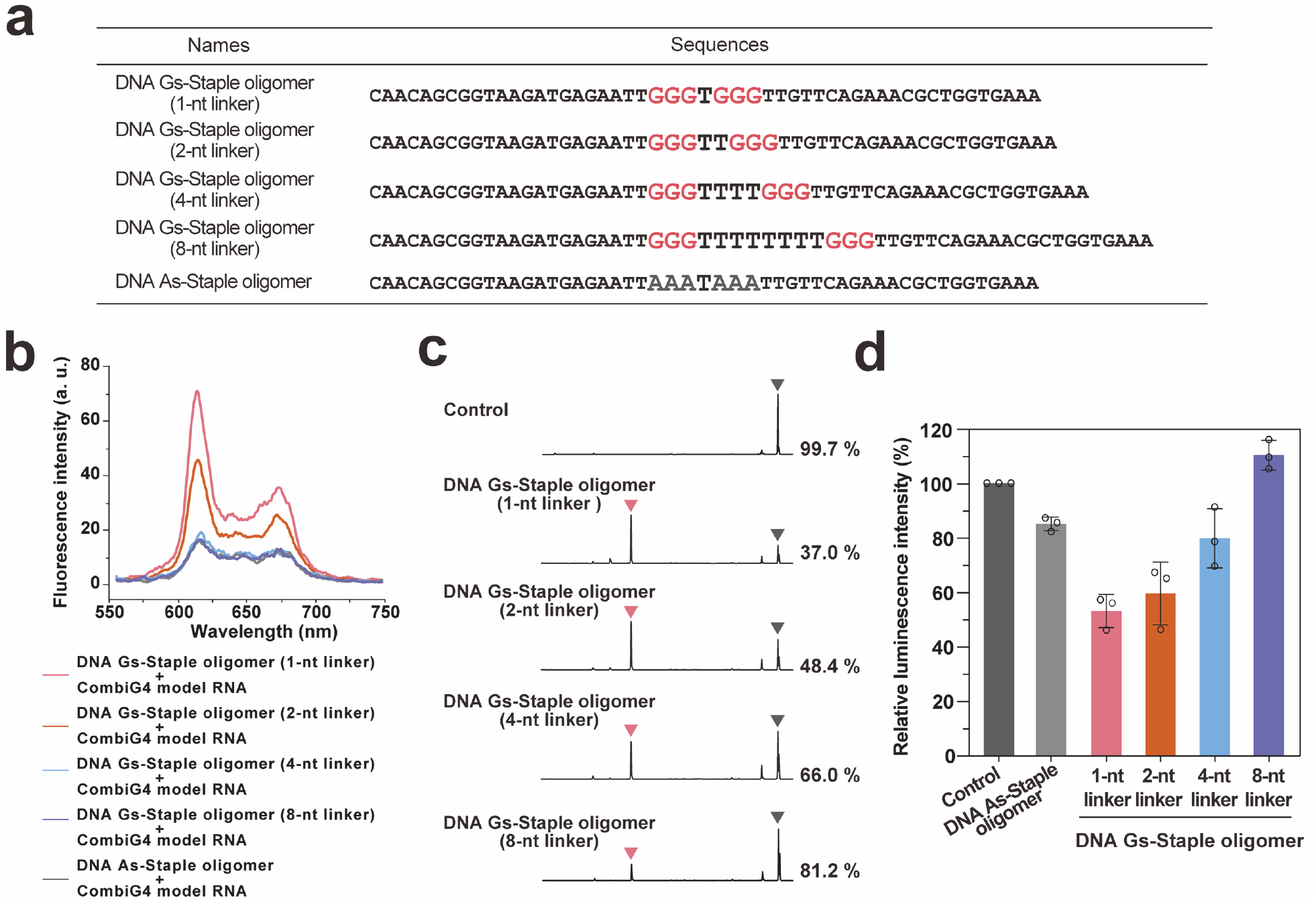
Characterizing the linker in Gs-Staple oligomer. **a,** Sequences of the DNA Gs-Staple oligomer and the DNA As-Staple oligomer. **b,** Evaluation of DNA/RNA combiG4 formation by fluorimetry (Ex. wavelength: 399 nm) with NMM. The curves show the fluorescence emission spectra of NMM in the presence of DNA Gs-Staple oligomer or As-Staple oligomer. **c,** RTase stop assay sequencing spectra of combiG4 model sequences. RTase-mediated cDNA synthesis was interrupted in the presence of all DNA Gs-Staple oligomers. Red arrowheads indicate the site of arrest of RTase with DNA/RNA combiG4 induced by the Gs-Staple oligomer, and black arrowheads indicate the end of RTase elongation without DNA/RNA combiG4 induction. The percentage represents the ratio of the full-length peak to the arrest site peak. **d,** Effect of the Gs-Staple oligomer on the translation of reporter RNA *in vitro*. Gs-Staple oligomer significantly reduced translation from reporter RNA templates. The translation suppression efficiency of Gs-Staple oligomers decreased with increasing length of the linker between the G-tracts of Gs-Staple oligomers. Data are the mean ± S.D. of three independent experiments.

The RTase stop assay was conducted to confirm the impact of linker length between G-tracts in DNA Gs-Staple oligomer. The results demonstrated that DNA Gs-Staple oligomer with the 1-nt linker showed the most effective inhibition of the progression of reverse transcription at the expected position and the efficacy of inhibition decreased with increasing linker length (Fig. 2c and S4c). Finally, we evaluated the effect of linker length on translation using an *in vitro* translation inhibition assay, employing a NanoLuc reporter gene that contained a 1+1_100-nt combiG4 model sequence in its 5’ UTR. An increase in the length of the linker resulted in a reduction in the efficiency of protein translation inhibition (Fig. 2d). In light of these findings, we selected the Gs-Staple oligomer with the 1-nt linker between G-tracts, which showed the highest translation inhibition effiicacy, for use in all subsequent experiments.

### Application of Gs-Staple oligomer to TRPC6 mRNAs

We introduced a Gs-Staple oligomer to the 5’ UTR of mouse TRPC6 (mTRPC6) mRNA, which is associated with cardiac hypertrophy (Fig. 3a and Table S2)^33, 34^. We first evaluated whether 31-, 41-, and 51-nt RNA-based Gs-Staple oligomers were able to induce combiG4 formation on the mTRPC6 sequence by measuring NMM fluorescence. Use of both the 41-nt and 51-nt RNA Gs-Staple oligomers resulted in a significant increase in fluorescence, whereas no signal change was observed with the 31-nt RNA Gs-Staple oligomer and the RNA As-Staple oligomer (Fig. 3b and Fig. S5a). The RTase stop assay demonstrated that the 41-nt RNA Gs-Staple oligomer effectively impedes the progression of reverse transcription (Fig. 3c and Fig. S5b). The introduction of the As-Staple oligomer did not produce peaks corresponding to the inhibition of reverse transcription at the expected combiG4 formation site. However, peaks indicative of inhibition of reverse transcription in the hybridization region were observed. This indicates that using RNA as the backbone nucleic acid in the As-Staple oligomer enhances binding affinity with the target RNA, thereby impeding reverse transcriptase progression in the hybridization region (Fig. 3c).

**Fig. 3.**
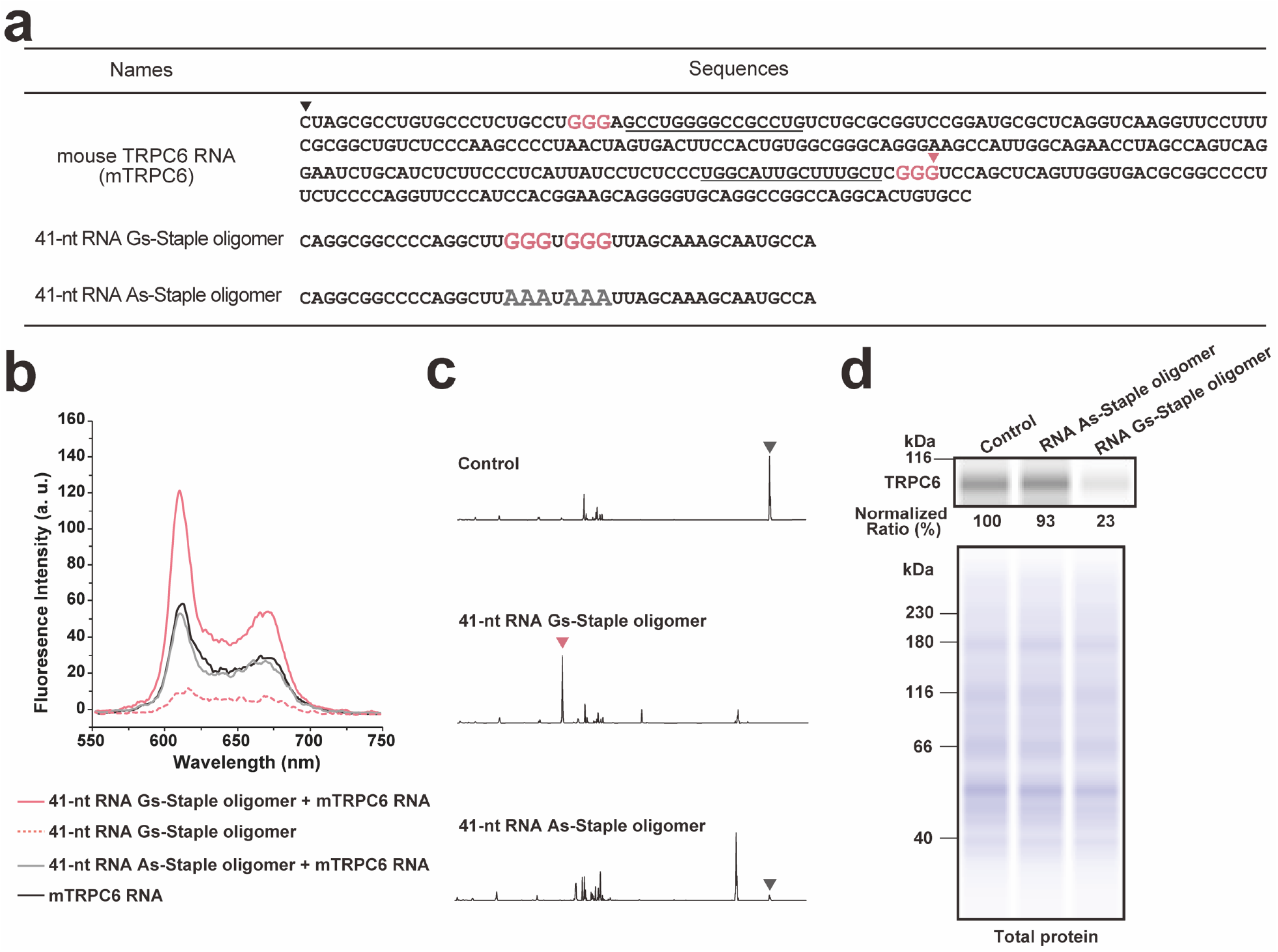
*In vitro* application of the Gs-Staple oligomer to the TRPC6 gene. **a,** Nucleotide sequences of the mouse TRPC6 mRNA. The G-tracts are shown in red. The red arrowhead indicates the site of arrest of RTase when a combiG4 structure is induced by the RNA Gs-Staple oligomer, and the black arrowhead indicates the end of RTase elongation without combiG4 induction. The underlines represent the 41-nt RNA Gs-or As-Staple oligomer recognition sites. **b,** Evaluation of combiG4 formation using NMM (Ex. wavelength: 399 nm). The red, dashed red, gray, and black curves show the curve shows the fluorescence emission spectra of NMM in the presence of 41-nt RNA Gs-Staple oligomer and mTRPC6 RNA, 41-nt RNA As-Staple oligomer alone, 41-nt RNA As-Staple oligomer and mTRPC6 RNA, and 41-nt RNA As-Staple oligomer alone, respectively. **c,** RTase stop assay of mTRPC6 RNA. RTase-mediated cDNA synthesis was interrupted on the 5’ UTR in the presence of the 41-nt RNA Gs-Staple oligomer. Red and gray arrowheads indicate the sites of arrest of RTase with combiG4 structures induced by the 41-nt RNA Gs-Staple oligomer and the end of RTase elongation without combiG4 influence, respectively. **d,** Evaluation of the effects of 41-nt RNA Gs-Staple oligomer on mTRPC6 expression in C2C12 cells by Western blotting (top image). *m*TRPC6 signals were normalized to total protein (bottom image).

Next, we evaluated the efficacy of the Gs-Staple oligomer in suppressing endogenous mTRPC6 protein expression in mammalian cells. The intracellular expression of the Gs-Staple oligomer was achieved via a short RNA expression vector, and the expression levels of the target proteins were subsequently assessed by Western blot analysis (Fig. S5c). The results demonstrated that C2C12 cells expressing Gs-Staple oligomer exhibited a substantial decrease in mTRPC6 protein expression levels. In contrast, C2C12 cells expressing As-Staple oligomer showed no discernible effect on protein expression (Fig. 3d). These findings provide strong evidence that the observed translation inhibition was due to the formation of a combiG4 between the G-tracts of Gs-Staple oligomer and those of the target mRNA. Furthermore, the observation that the expression levels of mTRPC6 remained unchanged in the presence of the As-Staple oligomer suggests that the hybridization of the RNA Gs-Staple oligomer to target mRNA does not lead to nonspecific changes in protein expression levels (Fig. 3d). These results suggest that, in addition to hybridization based on sequence complementarity, the formation of G4 structures is crucial for the effective function of Staples. This dual requirement is expected to assist in the minimization of off-target effects within the RNAh technology.

Subsequently, the Gs-Staple oligomer was applied to human TRPC6 (hTRPC6) mRNA to assess their versatility (Fig. S5a). We designed a DNA Gs-Staple oligomer that was specific to the hTRPC6 mRNA sequence and conducted a RTase stop assay. RTase stop assay results demonstrated that Gs-Staple oligomer effectively blocked the progression of reverse transcription at the DNA/RNA combiG4 formation site (Fig. S6b). Furthermore, HEK293T cells expressing the Gs-Staple oligomer showed a decrease in hTRPC6 protein expression levels compared to the control cells (Fig. S6c). These results reveal that the Gs-Staple oligomer is able to suppress endogenous protein expression levels.

### *In vivo* application of Gs-Staple oligomer

The functionality of the Gs-Staple oligomer was further evaluated *in vivo*. This experiment was conducted by introducing a short RNA expression vector encoding either the 41-nt RNA Gs-Staple oligomer or the 41-nt As-Staple oligomer into adeno-associated virus 6 (AAV6) (Fig. 4a)^35^. The AAV6 constructs were administered to mice via a tail vein injection. Two weeks after the injection, changes in mTRPC6 mRNA and protein expression levels in mouse heart tissue were evaluated by quantitative polymerase chain reaction (qPCR) and Western blot analysis. qPCR analysis revealed that there were no significant differences in the levels of mTRPC6 mRNA (Fig. 4b). In contrast, hearts expressing the RNA Gs-Staple oligomer showed a decrease in mTRPC6 protein levels, whereas those expressing the RNA As-Staple oligomer exhibited no discernible effect on protein expression (Fig. 4c). These results indicate that RNA Gs-Staple oligomer forms a combiG4 structure and suppresses protein translation processes without cleaving the target mRNA *in vivo*. Here, the Gs-Staple oligomer was shown to function as a RNAh tool with comparable performance to existing Staple oligomers that effectively suppresses the expression of a target protein that was previously inaccessible to the technology.

**Fig. 4.**
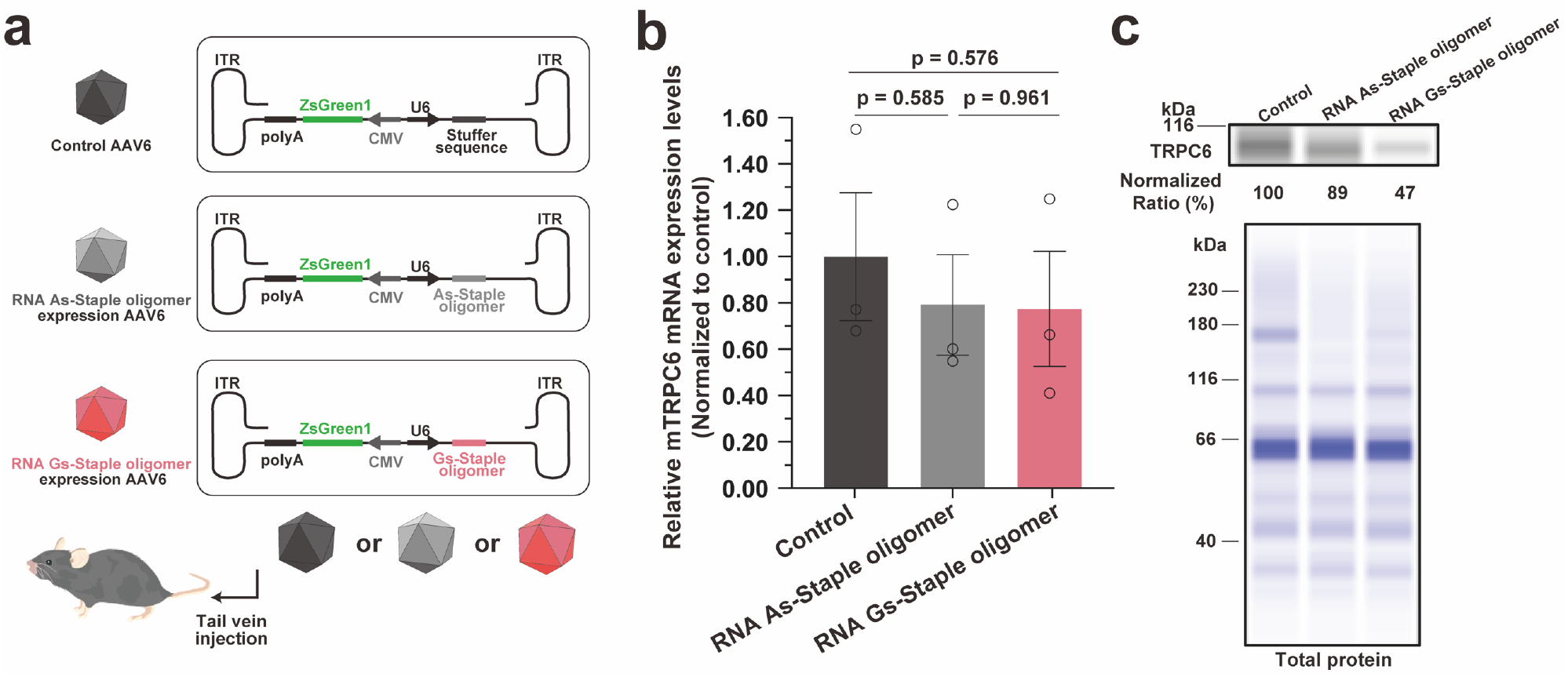
Effect of the Gs-Staple oligomer on *m*TRPC6 mRNA expression in mice. **a,** Schematic of the recombinant AAV6 vector genome encoding the Gs-Staple oligomer. A short hairpin RNA expression vector was used to express the Gs-Staple oligomer. The U6 promoter (black), Gs-Staple oligomer (red), As-Staple oligomer (gray) and Stuffer sequence (black) are shown, respectively. AAV6 infection is confirmed by ZsGreen (green) expression with the CMV promoter. The AAV6 expression system expresses the Gs-Staple oligomer in target organs. **b,** Evaluation of TRPC6 mRNA expression levels in each organ by qPCR. No significant change in the mRNA expression levels was observed in each organ by the introduction of the RNA Gs-Staple oligomer, clearly supporting the mechanism of action of RNAh that can suppress TRPC6 expression without mRNA degradation. Data are shown as the mean ± S.E. (n=3 mice/group, biologically independent samples). Student’s t-tests determined statistical significance. **c,** Evaluation of protein expression levels of TRPC6 in the heart by Western blotting (upper image). TRPC6 signals were normalized

## Discussion

Here, we developed the G-tract-supply Staple oligomer, Gs-Staple oligomer, representing a major advancement in RNAh technology by expanding its range of application. The Gs-Staple oligomer is characterized by the incorporation of G-tracts within its sequence. The design of the Staple oligomer enables it to hybridize with two distant regions on the target mRNA, facilitating the formation of a combiG4 structure in cooperation with the G-tracts in the target mRNA (Fig. 1b). This function overcomes the sequence constraints of conventional Staple oligomers, expanding the pool of target mRNAs. We have demonstrated that Gs-Staple oligomers effectively suppress gene expression by targeting hTRPC6 mRNA, which was difficult to achieve with the conventional Staple oligomer (Fig. S3b). Additionally, the relaxation of target sequence restrictions will enable the design of multiple Gs-Staple oligomers for a single mRNA. To further investigate the functionality of the Gs-Staple oligomer, we designed and evaluated multiple RNA-based Gs-Staple oligomers targeting mTRPC6 (Fig. S6a and Table S2). The results of NMM and RTase stop assays revealed that the NMM signal intensity and reverse transcription inhibition efficiency varied depending on the Gs-Staple oligomer sequence. Since mRNAs may form unique secondary structures depending on their sequences, some Staple oligomers may not have access to the targeted mRNA sequences, which may result in variations in their silencing efficacies. Even so, all of the Gs-Staple oligomers tested were shown to be functional *in vitro* (Fig. S6b-c). The increased flexibility provided by the ability to design multiple oligomers for a single gene makes sequence optimization for the Staple oligomer easier, and as a result, makes it possible to further minimize off-target effects and increase therapeutic efficacy compared to conventional Staple oligomers.

In this study, T or U linkers were incorporated between the G-tracts of the Gs-Staple oligomer. It was ascertained that an increasing number of linker nucleotides led to a decrease in the stability of the complex formed with the target RNA (Fig. S4b). Thus, the efficiency of both reverse transcription and translation inhibition decreased with increasing linker length (Fig. 2c-d and S4c). This observation aligns with previous studies that have demonstrated the importance of loop length between G-tracts in influencing the stability of G4 structures^36-38^. The findings of this study demonstrate that altering the length of the linker in the Gs-Staple oligomer similarly affects the stability of the induced combiG4 structure, thereby offering a means to modulate the potency of the Staple oligomer. The silencing activities of siRNAs and antisense oligonucleotides (ASOs) are contingent on the intracellular concentration of Argonaute proteins and RNaseH, whereas Gs-Staple oligomers function independently of endogenous enzymatic mechanisms. Consequently, the efficacy of gene silencing can be precisely regulated through rational molecular design. This control makes it a powerful and adaptable tool for functional genomics and the elucidation of gene regulatory mechanisms.

A significant aim in nucleic acid-based therapeutics, including RNA technologies, is to enhance *in vivo* stability and achieve targeted delivery to specific organs^39^. In this study, we employed AAV6-mediated organ targeting to achieve high-level expression of Gs-Staple oligomer *in vivo*, exploiting the infection specificity of AAV6 for precise delivery to target organs or cells. Unlike siRNAs and ASOs, Gs-Staple oligomers suppress protein expression through a distinct mechanism that does not depend on endogenous enzymes, allowing it to maintain efficacy even after chemical modification to reduce toxicity. In future research, it will be essential to introduce chemical modifications to Gs-Staple oligomer components for use as nucleic acid pharmaceuticals.

## Methods Oligonucleotides

All oligonucleotides were purchased from Thermo Fisher Scientific or Eurofin Genomics. DNA oligonucleotides were used as Gs-Staple oligomers, primers and PCR templates, and qPCR primers. RNA oligonucleotides were used as Gs-Staple oligomers. Individual DNA and RNA oligonucleotides were stored at a concentration of 100 µM in TE buffer.

### N-Methylmesoporphyrin IX (NMM) Fluorescence Measurements

A mixture of template RNA (0.24 µM) and staple oligomer (3.00 µM) in a buffer (20 mM Tris-HCl pH 7.8 and 100 mM KCl) was heated to 90°C for 2 min, and then gradually cooled to 20°C at a rate of 1.0°C/min. For measurement, NMM (GlpBio) was added at 0.1 µM to the mixture which was incubated for 30 min in the dark at room temperature. Fluorescence intensity was measured with a Cary Eclipse fluorescence spectrophotometer (Agilent Technologies). The excitation wavelength was at 399 nm. The fluorescence spectra were obtained by taking the moving average of five points made at 1.0 nm intervals from 550 to 750 nm.

### Thermal Denaturation Profiles

Melting experiments were conducted with target RNA and 5′-FAM-labeled staple oligomer hybrids (template: 0.3 μM, Staple: 0.5 μM) in 20 mM Tris-HCl buffer (pH 7.8) containing 150 mM KCl. The fluorescence intensity of the samples was monitored from 20 to 85°C with a heating rate of 0.5°C/min with the CFX Connect Real-Time System (Bio-Rad). The RNA-Staple oligomer hybrids were heated to 90°C and subsequently cooled to ambient temperature prior to measurements. The melting curves were converted to distinct melting peaks by plotting the first negative derivative of the fluorescence as a function of temperature (−dF/dT).

### RTase stop assay

A reaction mixture of template RNA (0.3 µM), Gs or As-Staple oligomer (1 µM), and a 5′-FAM-labeled primer [5′-FAM-CGC CAG GGT TTT CCC AGT CAC GAC-3′] (0.1 µM) was heated to 90°C for 2 min in folding buffer (50 mM Tris-HCl pH 7.8, containing 150 mM KCl and polyethylene glycol at 6.7 wt%), and then cooled to 20°C at a rate of 1.0°C/min. ReverTra Ace reverse transcriptase (Toyobo), MgCl_2_ (5 mM), and dNTPs (1.0 mM) were then added to the reaction mixture, and the reaction was carried out at 42 °C for 30 min, after which the reaction was stopped by heating at 99°C for 5 min. The produced cDNAs were analysed on a SeqStudio Genetic Analyzer (Applied Biosystems).

### *In Vitro* Translation Assay

A reaction mixture of template RNA (0.1 µM), with or without Gs-Staple oligomer or As-Staple oligomer (0.6 µM), and a buffer (50 mM Tris-HCl pH 7.8, 150 mM KCl) was heated to 90°C for 2 min, and then gradually cooled to 20°C at a rate of 1.0°C/min. The samples were used as RNA templates in 25 μL of cell-free protein expression mixture (Rabbit Reticulocyte Lysate), with or without Staple oligomer for 90 min at 30°C. Luciferase activity was evaluated using the Luciferase Assay kit (Promega) and a GloMax 20/20 Luminometer (Promega).

### Cell Cultures and Transfection

C2C12 and HEK293T cells were maintained in medium A (Dulbecco’s modified Eagle’s medium, supplemented with 100-units/mL penicillin, 100-mg/mL streptomycin sulfate, and 10% (v/v) fetal bovine serum). All cell lines were cultured at 37°C in a humidified 5% CO_2_ incubator. Each Staple-oligomer-expression vector was transfected into the cells using FuGENE HD (Promega) according to the manufacturer’s instructions.

### Animal Studies

Male C57BL/6J (7-week-old) mice were purchased from Japan SLC, Inc. All mice were maintained under a 12:12-h light–dark cycle, and the temperature was kept at 24°C with ad libitum access to a regular chow diet and water. All animal procedures were performed in accordance with Kumamoto University animal care guidelines and the Guide for the Care and Use of Laboratory Animals published by the U.S. National Institutes of Health (Publication No. 85-23, revised 1996) and permitted by the Animal Care and Use Committee of Kumamoto University.

### Tissue Collection and Histology

The heart and body weights of the sacrificed mice were recorded at the terminal point of the experiment. Portions of the myocardium were frozen in liquid nitrogen and stored at -80°C until use for Western blotting and RT-qPCR analysis.

### Adeno-associated Virus (AAV) Generation and Mouse Transduction

pAAV-U6-ZsGreen1 vector (5 μg), Staple oligomer-expression construct (5 μg), pRC6 vector (5 μg) and pHelper vector (5 μg) were mixed for co-transfection of HEK293T cells in Optimem medium (Thermo Fisher Scientific) using TransIT-VirusGEN^®^ Transfection Reagent (Mirus). 2 days after transfection, Staple-oligomer-expression or control AAVs were extracted from the cells and purified using AAVpro^®^ Purification Kit Maxi (All Serotype) (Takara Bio) according to the manufacturer’s protocol. The purified AAV titers were determined using AAVpro^®^ Titration Kit Ver.2 (Takara Bio) for real-time PCR according to the manufacturer’s protocol. The mice were anesthetized with isoflurane (Pfizer) for tail vein injections of AAVs. C57BL/6J mice were injected in the tail vein with 1.5×10^10^ vg of AAVs in 450 μL of PBS.

### Western Blot Analysis using the Abby Protein Simple System

In the cell translation assay, the cells were washed three times with cold PBS and lysed with RIPA buffer (Nacalai Tesque) containing a protease inhibitor cocktail (Nacalai Tesque). The cell lysates were passed through a 25G needle 10 times and centrifuged at 4°C for 10 min. In the *in vivo* translation assay, the mouse heart was homogenized with a lysis buffer (Cell Signaling Technology) containing a protease inhibitor cocktail (Nacalai Tesque) by using μT-12 bead beater homogenizer (TITEC) and SLPe40 ultrasonic homogenizer (BRANSON). The lysates were centrifuged at 4°C for 15 min to remove debris. The supernatants were transferred to new tubes and mixed with 6× sodium dodecyl sulfate (SDS) sample buffer (Nacalai Tesque), and then the mixture was heated at 95°C for 5 min. According to the manufacturer’s protocol, a simple Western blot was performed in an Abby instrument (Protein Simple) using 12–230kDa Separation 8×25 Capillary Cartridges (Protein Simple, SM-W004). The TRPC6 peak was detected by anti-TRPC6 antibody (Alomone Labs, ACC-017) used at a 1:50 dilution and Anti-Rabbit Detection Module (Protein Simple, DM-001). A total protein assay using the Total Protein Detection Module (Protein Simple, DM-TP01) and the Replex Module (Protein Simple, RP-001) was also performed in the same run. TRPC6 peak area was determined using Compass software (Protein Simple) and normalized to that of total protein.

### RT-qPCR

According to the manufacturer’s protocol, total RNA was isolated from cells or the mouse heart with ISOGEN (Nippon Gene). First-strand cDNAs were prepared by reverse transcription with an oligo (dT) primer (Thermo Fisher Scientific) and ReverTra Ace reverse transcriptase (TOYOBO) according to the manufacturer’s protocol. The cDNAs were subjected to qPCR, using a pair of GAPDH primers to quantify mouse GAPDH mRNA [5′-AAC AGC AAC TCC CAC TCT TCC-3′ and 5′-GTG GTC CAG GGT TTC TTA CTC -3′] and a pair of TRPC6 primers to quantify mouse TRPC6 mRNA [5′-AAC TCG GGG AGA GAC TG-3′ and 5′-ATA TGG CTT CAA GTG GAG-3′]. The qPCR analysis was performed on a MiniOpticon^TM^ Real-Time PCR System (Bio-Rad Laboratories) with SYBR® Green (Toyobo).

### Statistics and reproducibility

Results of representative experiments, such as Western blot analyses, were repeated at least two times independently with similar results. The figure legends provide the exact statistical test, P value, and sample size of each figure. The level of statistical significance is indicated by asterisks (\**P*<0.05; \*\**P*<0.01; \*\*\**P*<0.001). All attempts at replication were successful, and where only one result is shown (i.e. western blots, etc.), the data selected for publication are representative of the respective experiments.

### Data availability

The main data supporting the results of this study are available in the manuscript and its Supplementary Information section. The mTRPC6 and hTRPC6 mRNA sequences are available from the National Center for Biotechnology (NCBI) Nucleotide database, via accession numbers NM_001282086 and NM_001278190, respectively. The raw and analyzed datasets generated during the study are too large to publicly share. Still, they can be made available for research purposes on reasonable request to the corresponding authors.

## Acknowledgments

This work was supported by JSPS (23KJ1773 to T. Kida, 20H02769 to T.I., 22K05326, and 25K08834 to M.H., 20H02859, 21K18214, and 23H02082 to S. Sato), AMED under Grant Numbers JP20ak0101116, JP23ak0101168, JP21lm0203004, JP22ym0126806, JP22ym0126813, JP20lm0203012, JP24fk0310518, and JP22fk0410055 to Y. Katsuda. The Naito Foundation (T.I.), TERUMO Foundation for Life Sciences and Arts (T.I.), ZE Research Program, IAE (ZE2025B-18, and ZE2024B-18 to M.H.), and JST FOREST Program (JPMJFR211L to Y. K.). The authors thank Megumi Takashima for editing a draft of this manuscript.

## Author contributions

T.I., M.H., S. S., and Y. K., conceived the project and designed all experiments. T.K., Y.H., M.K., M.O., A.M., R.T., R.O., M.H., and Y.K., performed the experiments for *in vitro* characterization of the technology. T. K., Y.H., M.K., M.O., A.M., R.T., S.S., and Y. K., performed the experiments relating to *in vitro* and *in vivo* applications. T. K., Y.H., M.K., S.S., and Y. K., performed mouse injections and other mouse-related procedures. T.I., M.H., S. S., and Y.K., supervised this research and experimental design. T.K., T.I., M.H., S.S., and Y.K., wrote the manuscript with input from all authors.

## Competing interests

Y. K., Y.H., T.I., are coinventors on provisional patent applications (JP, no. 2022-81914) filed by Kumamoto University. Y. K., is a chief scientific officer and cofounder of StapleBio Inc. The other authors declare no competing interests.

