## Supplementary information for "Expanded gene targeting in RNA hacking with G-tract-supply Staple oligomer"

;

**Contents**

Supplementary Figure 1 | Evaluation of combiG4 formation using NMM fluorescent probe. ....................3
Supplementary Figure 2 | Identification of combiG4 formation on the combiG4 model sequences by RTase stop assay..........................................................................................................................4
Supplementary Figure 3 | Characterization of Gs-Staple oligomer to 2+1 type combiG4 model sequence. ..............................................................................................................................................5-6
Supplementary Method ...........................................................................................................13-15

**a**

| Names | Sequences |
| --- | --- |
| CombiG4 model seuqnece (1+1-60-nt) | –GGGUAUUCUCAUCUUACCGCUGUU- 60 nt -UUCACCAGCGUUUCUGAACCUGGG– |
| CombiG4 model seuqnece (1+1-100-nt) | –GGGUAUUCUCAUCUUACCGCUGUU-100 nt-UUCACCAGCGUUUCUGAACCUGGG– |
| CoombiG4 model seuqnece (1+1-140-nt) | –GGGUAUUCUCAUCUUACCGCUGUU-140 nt-UUCACCAGCGUUUCUGAACCUGGG– |
| CombiG4 model seuqnece (1+1-200-nt) | –GGGUAUUCUCAUCUUACCGCUGUU-200 nt-UUCACCAGCGUUUCUGAACCUGGG– |
| DNA Gs Staple oligomer (1+1 type) | AACAGCGGTAAGATGAGAATTTGGGTGGGTTGGTTCAGAAACGCTGGTGAA |
| DNA As Staple oligomer (1+1 type) | AACAGCGGTAAGATGAGAATTTAAATAAATTGGTTCAGAAACGCTGGTGAA |

**b**

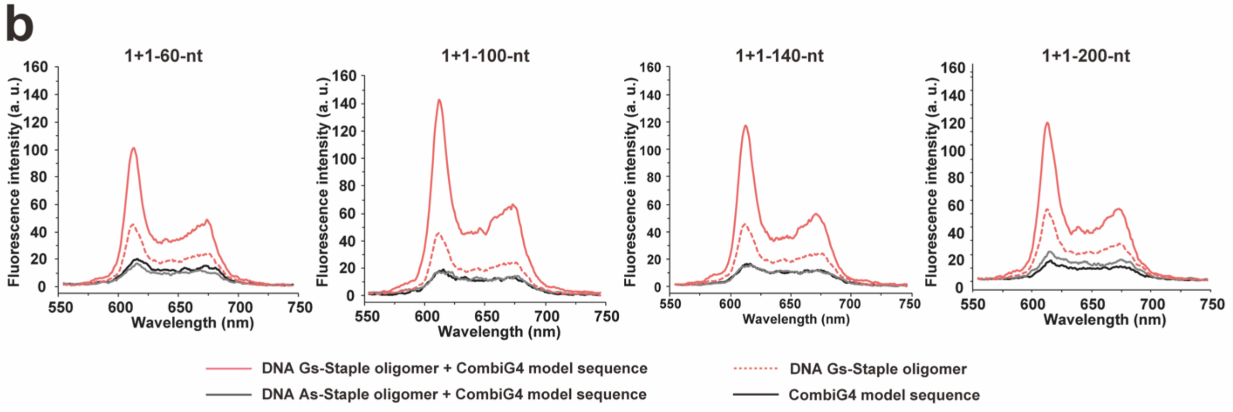

**Supplementary Figure 1 | Evaluation of combiG4 formation using NMM fluorescent probe. a,** The sequences of 1+1 type combiG4 models, DNA Gs-Staple oligomer and DNA As-Staple oligomer. The G-tracts are shown in red. The solid underlines represent the DNA Gs or DNA As-Staple oligomer recognition sites. **b,** Evaluation of DNA/RNA combiG4 formation by Gs-Staple oligomer with NMM. The red, gray and black curves show fluorescence emission spectra of NMM in the presence of DNA Gs-Staple oligomer and combiG4 model sequence RNA, As-Staple oligomer and combiG4 model sequence or combiG4 model sequence RNA only, respectively. The dashed curves represent the fluorescence emission spectra of NMM in the presence of DNA Gs-Staple oligomer.

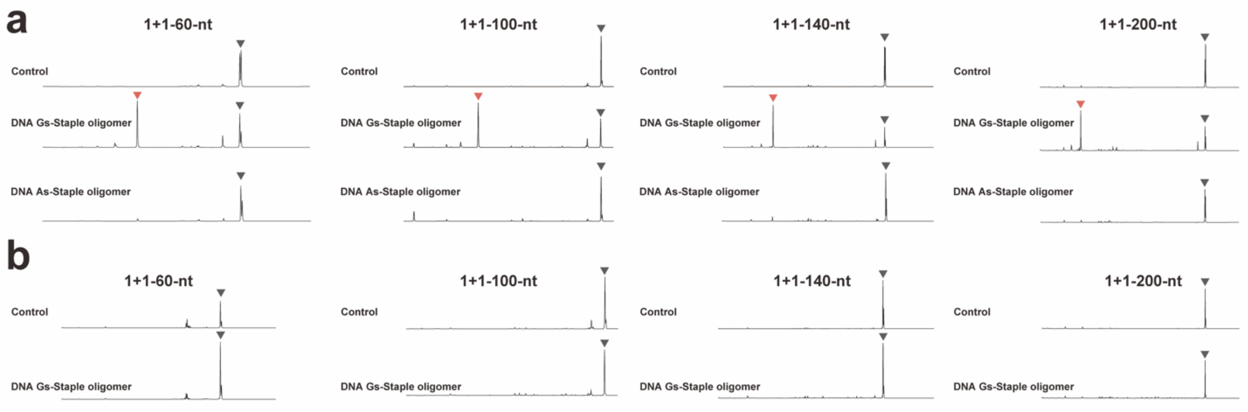

**Supplementary Figure 2 | Identification of combiG4 formation on the combiG4 model sequences by RTase stop assay. a,** Identification of sequencing spectra on 1+1 type combiG4 model sequences in the presence of KCl. RTase-mediated cDNA synthesis was interrupted on all 1+1 type combiG4 model sequences in the presence of Gs-Staple oligomer. Red and gray arrowheads indicate the sites of arrest of RTase with combiG4 induced by Gs-Staple oligomer and RTase elongation ends without combiG4 induction, respectively. **b,** Identification of sequencing spectra on 1+1 type combiG4 model sequences in the presence of LiCl. RTase-mediated cDNA synthesis in the presence of LiCl resulted in the elongation of cDNA sequences to their end. Gray arrowheads indicate RTase elongation end without combiG4 induction.

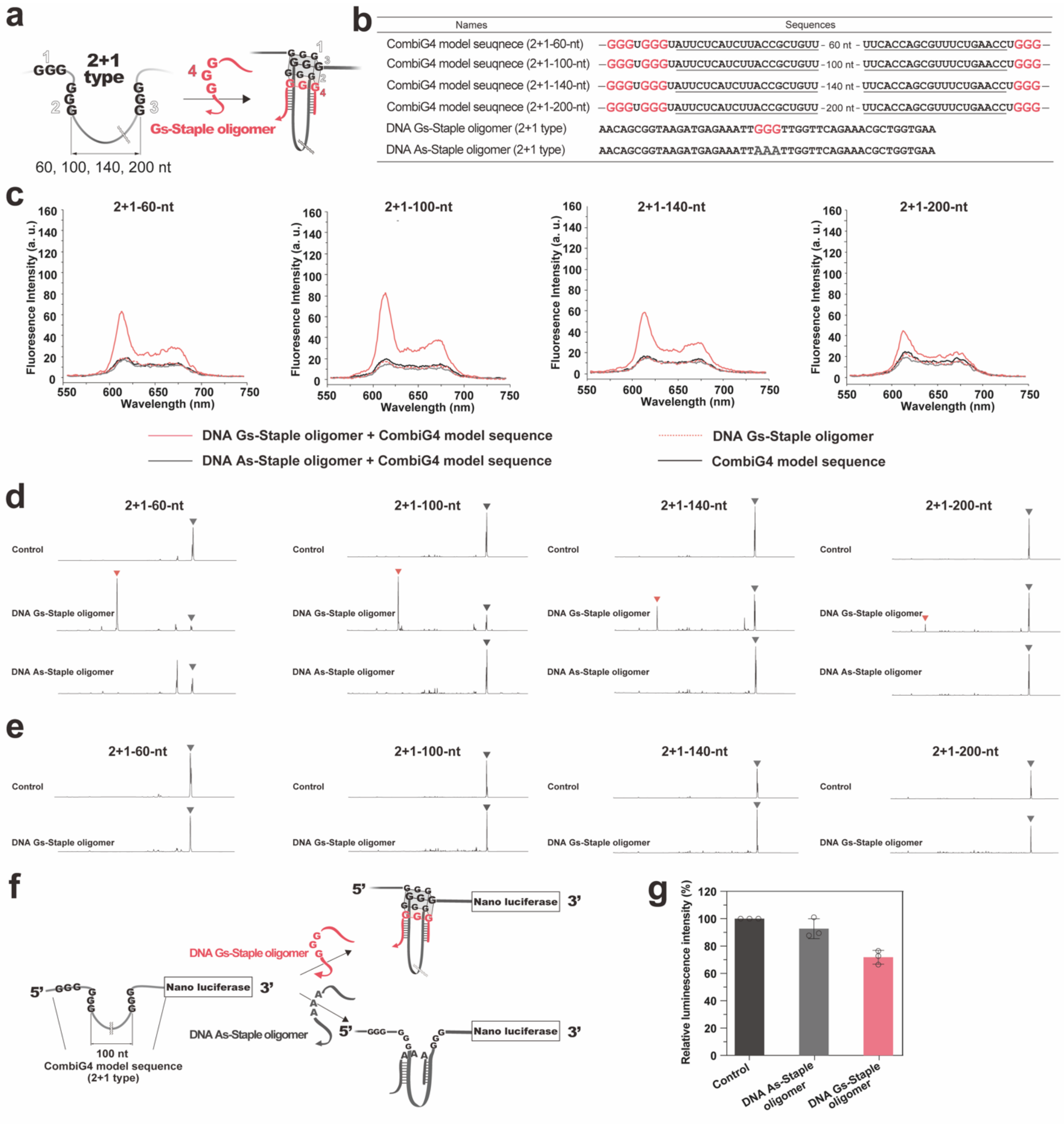

**Supplementary Figure 3 | Characterization of Gs-Staple oligomer to 2+1 type combiG4 model sequence. a,** 2+1 type combiG4 model sequences are illustrated. 2+1 type combiG4 model sequences have three G-tracts that are split into a 2+1 arrangement by 60-, 100-, 140- or 200-nt loops. **b,** The sequences of 2+1-type combiG4 model sequence, DNA Gs-Staple oligomer and DNA As-Staple oligomer. The G-tracts are shown in red. The solid underlines represent the DNA Gs-Staple oligomer or DNA As-Staple oligomer recognition sites. **c,** Evaluation of DNA/RNA combiG4 formation by Gs-Staple oligomer with NMM. The red, gray and black curves show fluorescence emission spectra of NMM in the presence of DNA Gs-Staple oligomer and combiG4 model sequence RNA, As-Staple oligomer and combiG4 model sequence or combiG4 model sequence RNA only, respectively. The dashed curves represent the fluorescence emission spectra of NMM in the presence of DNA Gs-Staple oligomer. **d,** Identification of spectra on combiG4 model sequence 1+1-100-nt by RTase stop assay. RTase-mediated cDNA synthesis was interrupted on combiG4 model sequences in the presence of each DNA Gs-Staple oligomer. Red arrowheads indicate arrest site of RTase with DNA/RNA combiG4 induced by the DNA Gs-Staple oligomer, and black arrowheads indicate the RTase elongation end without DNA/RNA combiG4 induction. **e,** Identification of sequencing spectra on 2+1 type combiG4 model sequences in the presence of LiCl. RTase-mediated cDNA synthesis in the presence of LiCl resulted in the elongation of cDNA sequences to their end. Gray arrowheads indicate RTase elongation end without combiG4 induction. **f,** 2+1-100-nt combiG4 model sequence was placed in the 5'UTR of Nano luciferase. The protein expression level was determined by Nanoluc luminescence intensity. DNA Gs-Staple oligomer induces DNA/RNA combiG4 formation (upper panel), whereas DNA As-Staple oligomer does not (lower panel). **g,** Effect of DNA Gs-Staple oligomer or As-Staple oligomer on the translation of reporter RNA *in vitro*. Data are mean ± S.D. of three independent experiments.

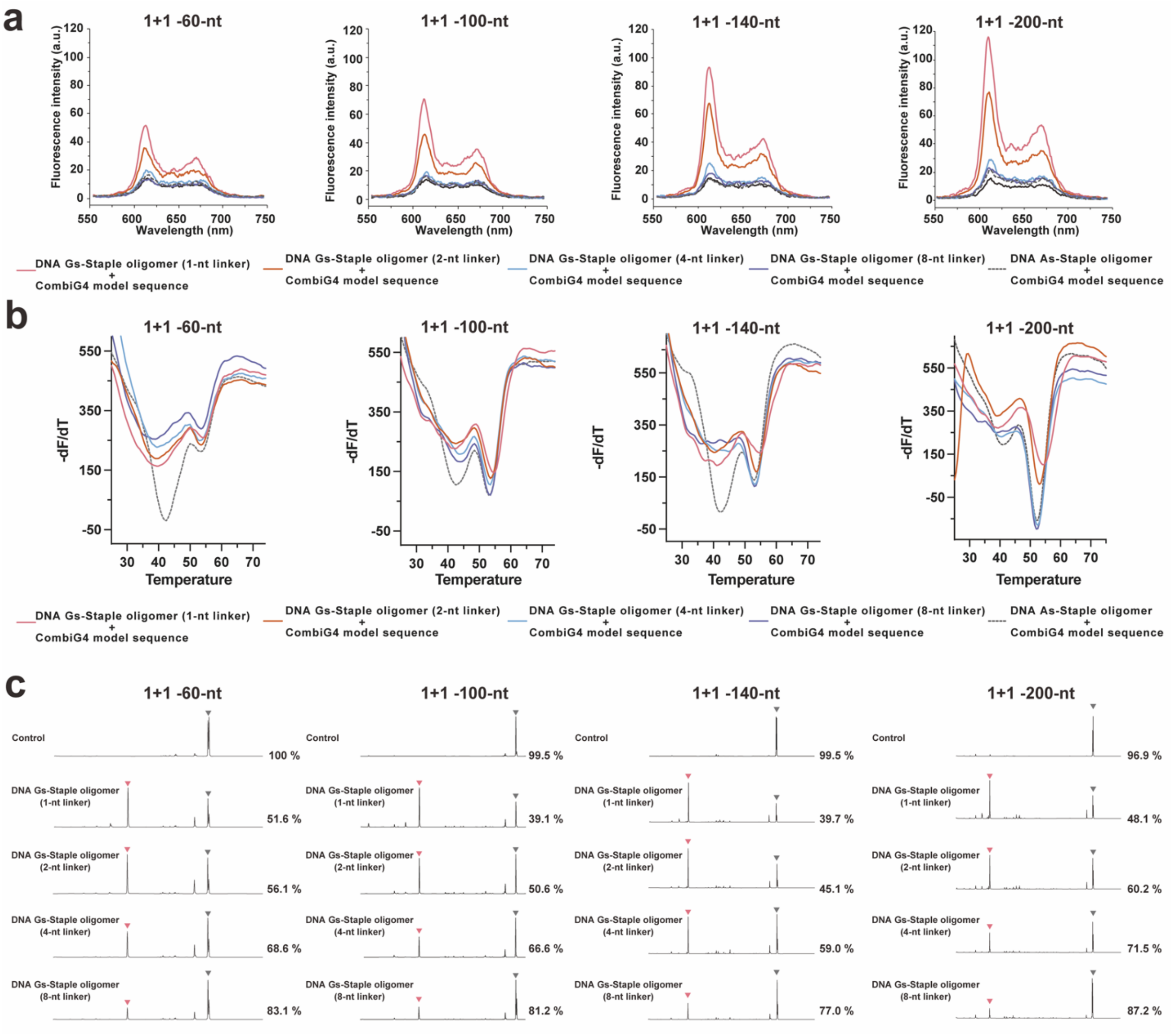

**Supplementary Figure 4 | Effect of the linker between G-tract in Gs-Staple oligomer *in vitro*. a,** Evaluation of combiG4 formation with NMM. **b,** Thermal denaturation profiles of Gs-Staple oligomer and combiG4 model sequence's RNA combis. The RNA strand containing either the combiG4 model sequence was complexed with a 5′-end FAM-labeled Gs or As-Staple oligomer, and the fluorescence intensity of the samples was measured from 20 to 85 °C. The graph displays the negative first derivative of the melting curve data (–dF/dT) versus temperature. **c,** Effect of the linker between G-tract in Gs-Staple oligomer on the RTase stop assay.

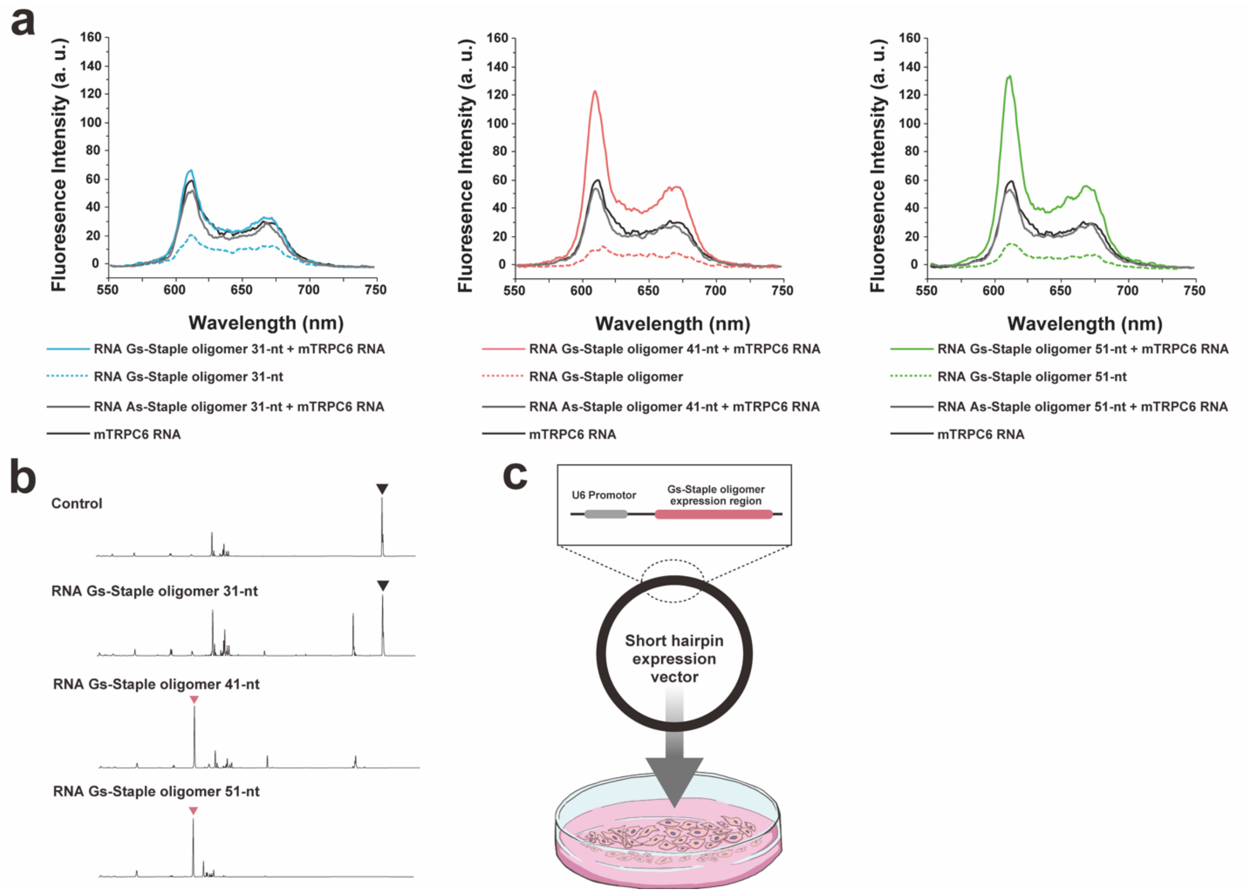

**Supplementary Figure 5 | *In vitro* and *in cell* application of Gs-Staple oligomer to the *m*TRPC6 mRNA. a,** Evaluation of combiG4 formation using NMM fluorescent probe. The red, green and blue curves show the fluorescence emission spectra of NMM in the presence of 51-nt, 41-nt or 31-nt RNA Gs-Staple oligomer, respectively. The solid and dashed curves represent the Gs-Staple oligomer and As-Staple oligomer, respectively. **b,** Identification of combiG4 formation on the 5'UTR of TRPC6 mRNA by RTase stop assay. RTase-mediated cDNA synthesis was interrupted on the 5'UTR in the presence of each RNA Staple oligomer. Red and black arrowheads indicate the sites of arrest of RTase with combiG4 induced by Staple oligomer and RTase elongation ends without combiG4 induction, respectively. **c,** Design and construction of an expression vector encoding Gs Staple oligomer. The short hairpin RNA expression vector was used to express an RNA Gs Staple oligomer. U6 promoter and Gs Staple oligomer are shown in gray and red, respectively.

**a**

| Names | Sequences |
|---|---|
| human TRPC6 mRNA | AUGAAGGCGGGAGCUGAGGGCUGGAGAGUCUCUGUUGACAUAGUAACUCUUCAGCUCCGUCUCC<br>CUUGCUCUCCGCUCUUACGCUUCGCUACCACCAGCGGCCCCGCCUGUGCCCUCUCUGCCCGGGC<br>GCCCCAGACGCAUCCUCGCGGGGUCUCCUCGGCCUGACCUGCUCAGGUCAAGAUCCUCUUUGC<br>ACCCCCUUAAGUGGUGACUUUUCCCCGGGCCAGUGGGCGAGCCACUUGCGGCGGGCGUCUGCAC<br>CCCCUGCUUCACCGUCGUCCCCUGGGCACCGGUCUGCCCAGGUCCAGUUCGGCCGCUGACGCGA<br>ACCCUCCGCACCGGGUCCCCGCUGGAACUGCCCACUCGGCUCCCCCGGGAGCGGGGCCCAGGC<br>CAGUCGGGCGUUCCCGC |
| DNA Gs Staple oligomer<br>(human TRPC6) | GAGCAGGTCAGGCCGAGGAGTTGGGGTGGGGTTCTCCCGGGGGAGCCGAGTGG |

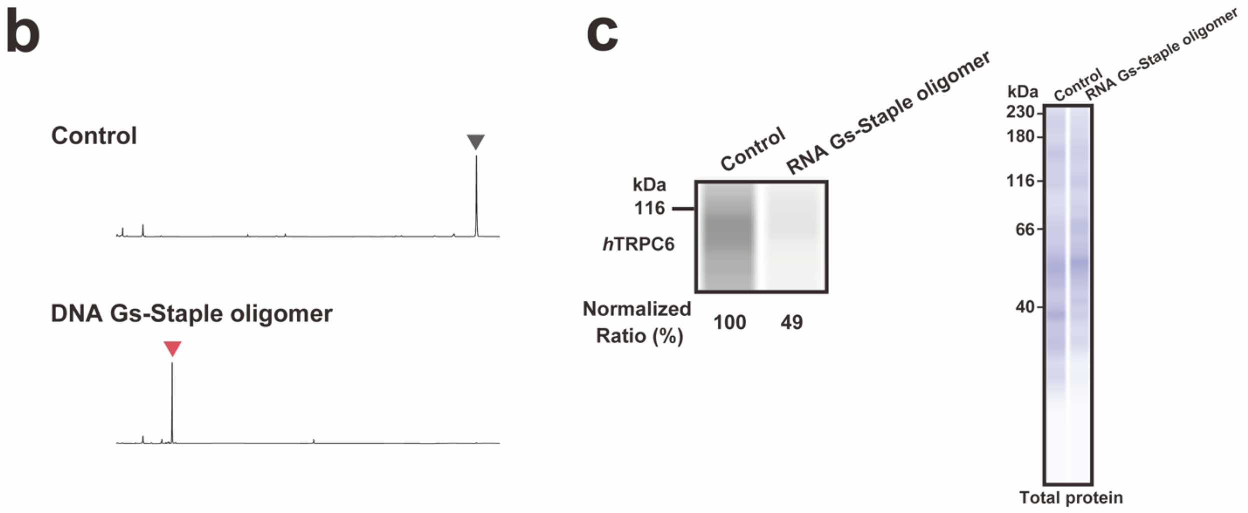

**Supplementary Figure 6 | *In vitro* and *in cell* application of Gs-Staple oligomer to the *h*TRPC6 mRNA.** **a,** Nucleotide sequences of the hTRPC6 and Gs-Staple oligomer. The G-tracts are shown in red. Underline shows a combiization site for the Gs-Staple oligomer. **b,** Identification of combiG4 formation on the hTRPC6 by RTase stop assay. RTase-mediated cDNA synthesis was interrupted on the 5' UTR in the presence of each Gs-Staple oligomer. Red and black arrowheads indicate the sites of arrest of RTase with combiG4 induced by Gs-Staple oligomer and RTase elongation ends without combiG4 induction, respectively. **c,** Evaluation of the effect of RNA Gs-Staple oligomers on hTRPC6 expression in HEK293T cells by western blotting. RNA Gs-Staple oligomer effectively suppressed hTRPC6 gene expression.

**a**

| | Names | Sequences |
|---|---|---|
| Design 1 | mouse TRPC6 RNA (mTRPC6) | ▼<br>CUAGCGCCUGUGCCCUCUGCCUGGGAGCCUGGGGCCGCCUGUCUGCGCGGUCCGGAUGCGCUCAGGUCAAGGUUCCUUU<br>CGCGGCUGUCUCCCAAGCCCCUAACUAGUGACUUCCACUGUGGCGGGCAGGGAAGCCAUUGGCAGAACCUAGCCAGUCA<br>GGAAUCUGCAUCUCUUCCCUCAUUAUCCUCUCCCUGGCAUUGCUUUGCUCGGGUCCAGCUCAGUUGGUGACGCGGCCCCU<br>UCUCCCCAGGUUCCCAUCCACGGAAGCAGGGGUGCAGGCCGGCCAGGCACUGUGCC |
| | RNA Gs-Staple oligomer 41-nt | CAGGCGGCCCCAGGCUUGGGUGGGUUCCACAGUGGAAGUCA |
| Design 2 | mouse TRPC6 RNA (mTRPC6) | ▼<br>CUAGCGCCUGUGCCCUCUGCCUGGGAGCCUGGGGCCGCCUGUCUGCGCGGUCCGGAUGCGCUCAGGUCAAGGUUCCUUUC<br>GCGGCUGUCUCCCAAGCCCCUAACUAGUGACUUCCACUGUGGCGGGCAGGGAAGCCAUUGGCAGAACCUAGCCAGUCAGG<br>AAUCUGCAUCUCUUCCCUCAUUAUCCUCUCCCUGGCAUUGCUUUGCUCGGGUCCAGCUCAGUUGGUGACGCGGCCCCUU<br>CUCCCCAGGUUCCCAUCCACGGAAGCAGGGGUGCAGGCCGGCCAGGCACUGUGCC |
| | RNA Gs-Staple oligomer 41-nt | CACCAACUGAGCUGGUUGGGUGGGUUGCUUCCGUGGAUGGG |
| Design 3 | mouse TRPC6 RNA (mTRPC6) | ▼<br>CUAGCGCCUGUGCCCUCUGCCUGGGAGCCUGGGGCCGCCUGUCUGCGCGGUCCGGAUGCGCUCAGGUCAAGGUUCCUUUC<br>GCGGCUGUCUCCCAAGCCCCUAACUAGUGACUUCCACUGUGGCGGGCAGGGAAGCCAUUGGCAGAACCUAGCCAGUCAG<br>GAAUCUGCAUCUCUUCCCUCAUUAUCCUCUCCCUGGCAUUGCUUUGCUCGGGUCCAGCUCAGUUGGUGACGCGGCCCCU<br>UCUCCCCAGGUUCCCAUCCACGGAAGCAGGGGUGCAGGCCGGCCAGGCACUGUGCC |
| | RNA Gs-Staple oligomer 41-nt | GCCAAUGGCUUCCCUUUGGGUGGGUUAGCAAAGCAAUGCCA |
| Design 4 | mouse TRPC6 RNA (mTRPC6) | ▼<br>CUAGCGCCUGUGCCCUCUGCCUGGGAGCCUGGGGCCGCCUGUCUGCGCGGUCCGGAUGCGCUCAGGUCAAGGUUCCUUUC<br>GCGGCUGUCUCCCAAGCCCCUAACUAGUGACUUCCACUGUGGCGGGCAGGGAAGCCAUUGGCAGAACCUAGCCAGUCAG<br>GAAUCUGCAUCUCUUCCCUCAUUAUCCUCUCCCUGGCAUUGCUUUGCUCGGGUCCAGCUCAGUUGGUGACGCGGCCCCUU<br>CUCCCCAGGUUCCCAUCCACGGAAGCAGGGGUGCAGGCCGGCCAGGCACUGUGCC |
| | RNA Gs-Staple oligomer 41-nt | GCCAATGGCUUCCCUUUGGGUGGGUUGCUUCCGUGGAUGGG |

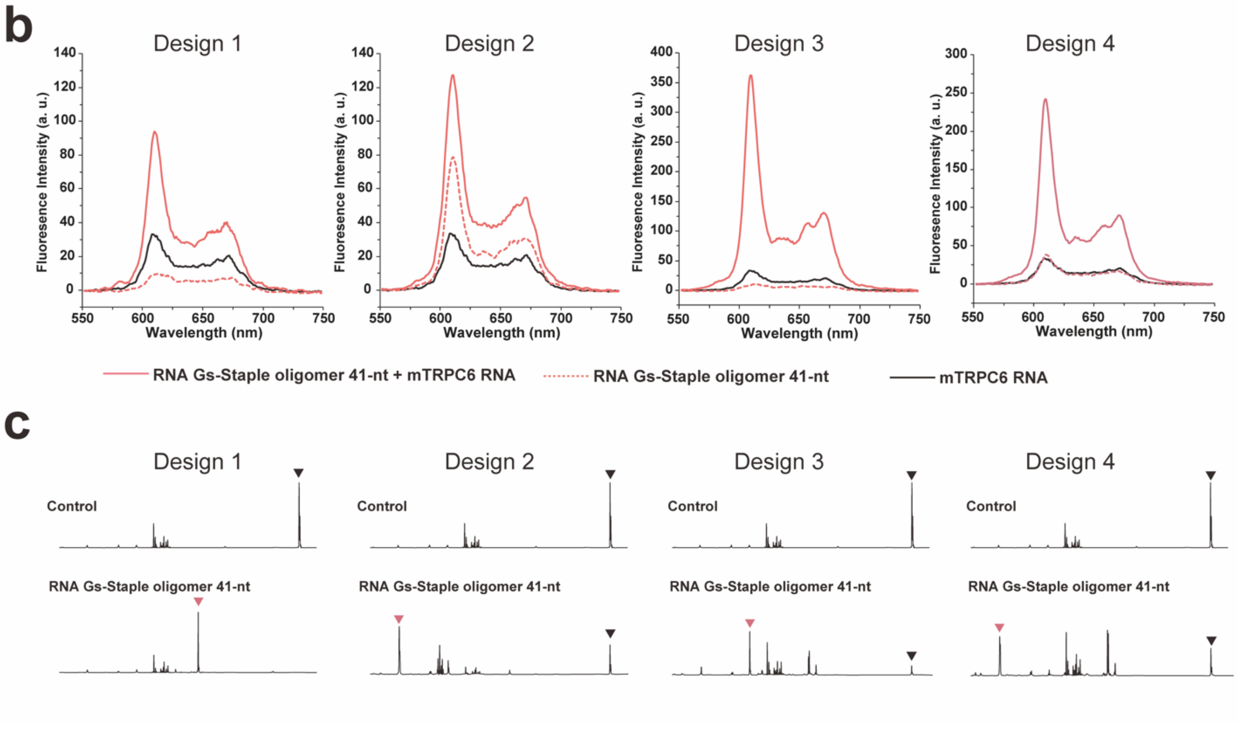

**Supplementary Figure 7 | *In vitro* evaluation of Gs-Staple oligomer against mTRPC6 gene. a,** Design of Gs-Staple oligomer. The G-tracts are shown in red. The solid underlines represent the RNA Gs-Staple oligomer recognition sites. **b,** Evaluation of combiG4 formation using NMM fluorescent probe. The red curves show the fluorescence emission spectra of NMM in the presence of RNA Gs-Staple oligomer 41-nt. **c,** Identification of combiG4 formation on the mTRPC6 RNA by RTase stop assay. RTase-mediated cDNA synthesis was interrupted on the 5’UTR in the presence of each RNA Gs-Staple oligomer. Red and gray arrowheads indicate the sites of arrest of RTase with combiG4 induced by RNA Gs-Staple oligomer 41-nt and RTase elongation ends without combiG4 induction, respectively.

**Supplementary Table 1 |** Nucleotide sequences of the target RNA and Nano luciferase RNA.

| Names | Sequences |
|---|---|
| **1+1-60-nt (CombiG4 model sequence)** | GAUCCUUUGAGGGUAUUCUCAUCUUACCGCUGUUGAGAUCCAGUCUUUUACUUUCACCAGCGUUUCUGAACCUGGGAUUCGCG<br>UUACCCGGGUACCGAGCUCGAAUUCACUGGCCGUCGUUUUACAACGUCGUGACUGGGAAAACCCUGGCG |
| **1+1-100-nt (CombiG4 model sequence)** | GAUCCUUUGAGGGUAUUCUCAUCUUACCGCUGUUGAGAUCCAGUUCGAUGUAACCCACUCGUGCACCCAACUGAUCUUCAGCA<br>UCUUUUACUUUCACCAGCGUUUCUGAACCUGGGAUUCGCGUUACCCGGGUACCGAGCUCGAAUUCACUGGCCGUCGUUUUACA<br>ACGUCGUGACUGGGAAAACCCUGGCG |
| **1+1-140-nt (CombiG4 model sequence)** | GAUCCUUUGAGGGUAUUCUCAUCUUACCGCUGUUGAGAUCCAGUUCGAUGUAACCCACUCGUGCGCAGGAAAGAACAUGUGAG<br>CAAAAGGCCAGCAAAAGGCCCACCCAACUGAUCUUCAGCAUCUUUUACUUUCACCAGCGUUUCUGAACCUGGGAUUCGCGUUA<br>CCCGGGUACCGAGCUCGAAUUCACUGGCCGUCGUUUUACAACGUCGUGACUGGGAAAACCCUGGCG |
| **1+1-200-nt (CombiG4 model sequence)** | CAGGUCGACUCUAGAGUAAUACGACUCACUAUAGAUCCUUUGAGGGUAUUCUCAUCUUACCGCUGUUGAGAUCCAGUUCGAUG<br>UAACCCACUCGUGCGCAGGAAAGAACAUGUGAGUCGACGCUCAAGUCAGAGGUGGCGAAACCCGACAGGACUAUAAAGAUACC<br>AGGCGUUUCCCAAAAGGCCAGCAAAAGGCCCACCCAACUGAUCUUCAGCAUCUUUUACUUUCACCAGCGUUUCUGAACCUGGG<br>AUUCGCGUUACCCGGGUACCGAGCUCGAAUUCACUGGCCGUCGUUUUACAACGUCGUGACUGGGAAAACCCUGGCG |
| **2+1-60-nt (CombiG4 model sequence)** | GAUCCUUUGAGGGUGGGUAUUCUCAUCUUACCGCUGUUGAGAUCCAGUCUUUUACUUUCACCAGCGUUUCUGAACCUGGGAUU<br>CGCGUUACCCGGGUACCGAGCUCGAAUUCACUGGCCGUCGUUUUACAACGUCGUGACUGGGAAAACCCUGGCG |
| **2+1-100-nt (CombiG4 model sequence)** | GAUCCUUUGAGGGUGGGUAUUCUCAUCUUACCGCUGUUGAGAUCCAGUUCGAUGUAACCCACUCGUGCACCCAACUGAUCUUC<br>AGCAUCUUUUACUUUCACCAGCGUUUCUGAACCUGGGAUUCGCGUUACCCGGGUACCGAGCUCGAAUUCACUGGCCGUCGUUU<br>UACAACGUCGUGACUGGGAAAACCCUGGCG |
| **2+1-140-nt (Binary G4 model sequence)** | GAUCCUUUGAGGGUGGGUAUUCUCAUCUUACCGCUGUUGAGAUCCAGUUCGAUGUAACCCACUCGUGCGCAGGAAAGAACAUG<br>UGAGCAAAAGGCCAGCAAAAGGCCCACCCAACUGAUCUUCAGCAUCUUUUACUUUCACCAGCGUUUCUGAACCUGGGAUUCGC<br>GUUACCCGGGUACCGAGCUCGAAUUCACUGGCCGUCGUUUUACAACGUCGUGACUGGGAAAACCCUGGCG |
| **2+1-200-nt (CombiG4 model sequence)** | CAGGUCGACUCUAGAGUAAUACGACUCACUAUAGAUCCUUUGAGGGUGGGUAUUCUCAUCUUACCGCUGUUGAGAUCCAGUUC<br>GAUGUAACCCACUCGUGCGCAGGAAAGAACAUGUGAGUCGACGCUCAAGUCAGAGGUGGCGAAACCCGACAGGACUAUAAAGA<br>UACCAGGCGUUUCCCAAAAGGCCAGCAAAAGGCCCACCCAACUGAUCUUCAGCAUCUUUUACUUUCACCAGCGUUUCUGAACC<br>UGGGAUUCGCGUUACCCGGGUACCGAGCUCGAAUUCACUGGCCGUCGUUUUACAACGUCGUGACUGGGAAAACCCUGGCG |
| **5' UTR of mouse TRPC6 mRNA** | CGCCUGUGCCCUCUGCCUGGGAGCCUGGGGCCGCCUGUCUGCGCGGUCCGGAUGCGCUCAGGUCAAGGUUCCUUUCGCGGCUG<br>UCUCCCAAGCCCCUAACUAGUGACUUCCACUGUGGCGGGCAGGGAAGCCAUUGGCAGAACCUAGCCAGUCAGGAAUCUGCAUC<br>UCUUCCCUCAUUAUCCUCUCCCUGGCAUUGCUUUGCUCGGGUCCUCCACGGAAGCAGGGUGCAGGCCGGCCAGGCACUGUGCC<br>AUG |
| **5' UTR human TRPC6 mRNA** | AUGAAGGCGGGAGCUGAGGGCUGGAGAGUCUCUGUUGACAUAGUAACUCUUCAGCUCCGUCUCCCUUGCUCUCCGCUCUUACG<br>CUUCGCUACCACCAGCGGCCCCGCCUGUGCCCUCUCUGCCCGGGCGCCCCAGACGCAUCCUCGCGGGGUCUCCUCGGCCUGAC<br>CUGCUCAGGUCAAGAUCCUCUUUGCACCCCCUUAAGUGGUGACUUUUCCCCGGGCCAGUGGGCGAGCCACUUGCGGCGGGCGU<br>CUGCACCCCCUGCUUCACCGUCGUCCCCUGGGCACCGGUCUGCCCAGGUCCAGUUCGGCCGCUGACGCGAACCCUCCGCACCG<br>GGUCCCCGCUGGAACUGCCCACUCGGCUCCCCCGGGAGCGGGGCCCAGGCCAGUCGGGCGUUCCCGC |
| **Nano luciferase RNA** | AUGGUCUUCACACUCGAAGAUUUCGUUGGGGACUGGCGACAGACAGCCGGCUACAACCUGGACCAAGUCCUUGAACAGGGAGG<br>UGUGUCCAGUUUGUUUCAGAAUCUCGGGGUGUCCGUAACUCCGAUCCAAAGGAUUGUCCUGAGCGGUGAAAAUGGGCUGAAGA<br>UCGACAUCCAUGUCAUCAUCCCGUAUGAAGGUCUGAGCGGCGACCAAAUGGGCCAGAUCGAAAAAAUUUUUAAGGUGGUGUAC<br>CCUGUGGAUGAUCAUCACUUUAAGGUGAUCCUGCACUAUGGCACACUGGUAAUCGACGGGGUUACGCCGAACAUGAUCGACUA<br>UUUCGGACGGCCGUAUGAAGGCAUCGCCGUGUUCGACGGCAAAAAGAUCACUGUAACAGGGACCCUGUGGAACGGCAACAAAA<br>UUAUCGACGAGCGCCUGAUCAACCCCGACGGCUCCCUGCUGUUCCGAGUAACCAUCAACGGAGUGACCGGCUGGCGGCUGUGC<br>GAACGCAUUCUGGCGUAA |

**Supplementary Table 2** | Nucleotide sequences of Gs or As-Staple oligomers for combiG4 model sequence, mTRPC6, and hTRPC6.

| Names | Sequences |
|---|---|
| DNA Gs-Staple oligomer 51-nt (1-nt linker, 1+1 type) | CAACAGCGGTAAGATGAGAATTGGGTGGGTTGTTCAGAAACGCTGGTGAAA |
| DNA Gs-Staple oligomer 52-nt (2-nt linker, 1+1 type) | CAACAGCGGTAAGATGAGAATTGGGTTGGGTTGTTCAGAAACGCTGGTGAAA |
| DNA Gs-Staple oligomer 54-nt (4-nt linker, 1+1 type) | CAACAGCGGTAAGATGAGAATTGGGTTTTGGGTTGTTCAGAAACGCTGGTGAAA |
| DNA Gs-Staple oligomer 58-nt (8-nt linker, 1+1 type) | CAACAGCGGTAAGATGAGAATTGGGTTTTTTTTGGGTTGTTCAGAAACGCTGGTGAAA |
| DNA As-Staple oligomer 51-nt (1+1 type) | CAACAGCGGTAAGATGAGAATTAAATAAATTGTTCAGAAACGCTGGTGAAA |
| DNA Gs-Staple oligomer 47-nt (2+1 type) | CAACAGCGGTAAGATGAGAATTGGGTTGTTCAGAAACGCTGGTGAAA |
| DNA As-Staple oligomer 47-nt (2+1 type) | CAACAGCGGTAAGATGAGAATTAAATTGTTCAGAAACGCTGGTGAAA |
| RNA Gs-Staple oligomer 31-nt (mTRPC6) | GGCCCCAGGCUUGGGUGGGUUAGCAAAGCAA |
| RNA Gs-Staple oligomer 41-nt (mTRPC6) | CAGGCGGCCCCAGGCUUGGGUGGGUUAGCAAAGCAAUGCCA |
| RNA As-Staple oligomer 41-nt (mTRPC6) | CAGGCGGCCCCAGGCUUAAAUAAAUUAGCAAAGCAAUGCCA |
| RNA Gs-Staple oligomer 51-nt (mTRPC6) | GCAGACAGGCGGCCCCAGGCUUGGGUGGGUUAGCAAAGCAAUGCCAGGGAG |
| RNA Gs-Staple oligomer 41-nt (Design 1, mTRPC6) | CAGGCGGCCCCAGGCUUGGGUGGGUUCCACAGUGGAAGUCA |
| RNA Gs-Staple oligomer 41-nt (Design 2, mTRPC6) | CACCAACUGAGCUGGUUGGGUGGGUUGCUUCCGUGGAUGGG |
| RNA Gs-Staple oligomer 41-nt (Design 3, mTRPC6) | GCCAAUGGCUUCCCUUUGGGUGGGUUAGCAAAGCAAUGCCAA |
| RNA Gs-Staple oligomer 41-nt (Design 4, mTRPC6) | GCCAATGGCUUCCCUUUGGGUGGGUUGCUUCCGUGGAUGGG |
| DNA Gs-Staple oligomer 53-nt (hTRPC6) | GAGCAGGTCAGGCCGAGGAGTTGGGGTGGGGTTCTCCCGGGGGAGCCGAGTGG |
| RNA Gs-Staple oligomer 53-nt (hTRPC6) | GAGCAGGUCAGGCCGAGGAGUUGGGGUGGGGUUCUCCCGGGGGAGCCGAGUGG |

**Supplementary Methods**

**Preparation of pUC19-1+1 type combiG4 model sequence Constructs**

**pUC19-1+1-60-nt combiG4 model sequence:**

The 1+1-60-nt DNA fragment was prepared by PCR using DNA [5′-CAG GTC GAC TCT AGA GTA ATA CGA CTC ACT ATA GAT CCT TTG AGG GTA TTC TCA TCT TAC CGC TGT TGA GAT CCA GTC TTT TAC TTT CAC CAG CGT TTC TGA ACC TGG GAT TCG CGT TAC CCG GGT ACC GAG CTC-3′] as a template with a forward [5′-CAG GTC GAC TCT AGA GTA ATA CGA CTC ACT ATA GAT CCT TTG AGG-3′] and a reverse primer [5′-GAG CTC GGT ACC CGG GTA ACG CGA ATC C-3′]. The PCR product was subcloned into the Bam HI site in pUC19 vector by using HiFi DNA Assembly Cloning Kit (New England Biolabs), then pUC19-1+1-60-nt combiG4 model sequence construct was obtained.

**pUC19-1+1-100-nt combiG4 model sequence:**

The 1+1-100-nt DNA fragment was prepared by PCR using pSuper vector as a template with a forward primer [5′-ATC TTA CCG CTG TTG AG-3′] and a reverse primer [5′-AGA AAC GCT GGT GAA AG-3′]. The dsDNA was subsequently amplified by PCR, using a forward primer [5′-TAT AGA TCC TTT GAG GGT ATT CTC ATC TTA CCG CTG TTG AG-3′] and a reverse primer [5′-CGG GTA ACG CGA ATC CCA GGT TCA GAA ACG CTG GTG AAA GT-3′]. The dsDNA was subsequently amplified by PCR, using a forward primer [5′-CAG GTC GAC TCT AGA GTA ATA CGA CTC ACT ATA GAT CCT TTG AGG-3′] and a reverse primer [5′-GAG CTC GGT ACC CGG GTA ACG CGA ATC C-3′]. The PCR product was subcloned into the Bam HI site in pUC19 vector by using HiFi DNA Assembly Cloning Kit (New England Biolabs), then pUC19-1+1-100-nt combiG4 model sequence construct was obtained.

**pUC19-1+1-140-nt combiG4 model sequence:**

The 1+1-140-nt DNA fragment was prepared by overlap extension PCR. Four dsDNA fragments were prepared by DNA polymerase reaction, using synthetic oligonucleotides [5′-CAG GTC GAC TCT AGA GTA ATA CGA CTC ACT ATA GAT CCT TTG AGG GTA TTC TCA TCT TAC CGC TGT TGA GAT CCA GTT CGA TGT AAC CCA CTC GTG CGC AGG AAA GAA CAT GTG AGC AAA AGG CC-3′] and [5′-GAG CTC GGT ACC CGG GTA ACG CGA ATC CCA GGT TCA GAA ACG CTG GTG AAA GTA AAA GAT GCT GAA GAT CAG TTG GGT GGG CCT TTT GCT GGC CTT TTG CTC ACA TGT TCT TTC CTG CGC-3′]. The dsDNA was subsequently amplified by PCR, using a forward primer [5′-CAG GTC GAC TCT AGA GTA ATA CGA CTC ACT ATA GAT CCT TTG AGG-3′] and a reverse primer [5′-GAG CTC GGT ACC CGG GTA ACG CGA ATC C-3′]. The PCR product was subcloned into the Bam HI site in pUC19 vector by using HiFi DNA Assembly Cloning Kit (New England Biolabs), then pUC19-1+1-140-nt combiG4 model sequence construct was obtained.

**pUC19-1+1-200-nt combiG4 model sequence:**

The 1+1-200nt DNA fragment was prepared by overlap extension PCR. Four dsDNA fragments were prepared by DNA polymerase reaction, using synthetic oligonucleotides [5′-GGG TAT TCT CAT CTT ACC GCT GTT GAG ATC CAG TTC GAT GTA ACC CAC TCG TGC GCA GGA AAG AAC ATG TGA GTC GAC GCT CAA GTC AGA GGT GGC GAA ACC CGA CAG GAC TAT AAA GAT ACC AGG CG-3′] and [5′-AGG TTC AGA AAC GCT GGT GAA AGT AAA AGA TGC TGA AGA TCA GTT GGG TGG GCC TTT TGC TGG CCT TTT GGG AAA CGC CTG GTA TCT TTA TAG TCC TGT CGG G-3′]. The dsDNA was subsequently amplified by PCR, using a forward primer [5′-TAT AGA TCC TTT GAG GGT ATT CTC ATC TTA CCG CTG TTG AG-3′] and a reverse primer [5′-CGG GTA ACG CGA ATC CCA GGT TCA GAA ACG CTG GTG AAA GT-3′]. The dsDNA was subsequently amplified by PCR, using a forward primer [5′-CAG GTC GAC TCT AGA GTA ATA CGA CTC ACT ATA GAT CCT TTG AGG-3′] and a reverse primer [5′-GAG CTC GGT ACC CGG GTA ACG CGA ATC C-3′]. The PCR product was subcloned into the Bam HI site in pUC19 vector by using HiFi DNA Assembly Cloning Kit (New England Biolabs), then pUC19-1+1-200-nt combiG4 model sequence construct was obtained.

**Preparation of pUC19-2+1 type combiG4 model sequence Constructs**

**pUC19-2+1-60-nt combiG4 model sequence:**

The 2+1-60-nt DNA fragment was prepared by PCR using DNA [5′-CAG GTC GAC TCT AGA GTA ATA CGA CTC ACT ATA GAT CCT TTG AGG GTG GGT ATT CTC ATC TTA CCG CTG TTG AGA TCC AGT CTT TTA CTT TCA CCA GCG TTT CTG AAC CTG GGA TTC GCG TTA CCC GGG TAC CGA GCT C-3′] as a template with a forward [5′-CAG GTC GAC TCT AGA GTA ATA CGA CTC ACT ATA GAT CCT TTG AGG-

3′] and a reverse primer [5′-GAG CTC GGT ACC CGG GTA ACG CGA ATC C-3′]. The PCR product was subcloned into the Bam HI site in pUC19 vector by using HiFi DNA Assembly Cloning Kit (New England Biolabs), then pUC19-2+1-60-nt combiG4 model sequence construct was obtained.

**pUC19-1+1-100-nt combiG4 model sequence:**

The 2+1-100-nt DNA fragment was prepared by PCR using pSuper vector as a template with a forward primer [5′-ATC TTA CCG CTG TTG AG-3′] and a reverse primer [5′-AGA AAC GCT GGT GAA AG-3′]. The dsDNA was subsequently amplified by PCR, using a forward primer [5′-TAT AGA TCC TTT GAG GGT GGG TAT TCT CAT CTT ACC GCT GTT GAG-3′] and a reverse primer [5′-CGG GTA ACG CGA ATC CCA GGT TCA GAA ACG CTG GTG AAA GT-3′]. The dsDNA was subsequently amplified by PCR, using a forward primer [5′-CAG GTC GAC TCT AGA GTA ATA CGA CTC ACT ATA GAT CCT TTG AGG-3′] and a reverse primer [5′-GAG CTC GGT ACC CGG GTA ACG CGA ATC C-3′]. The PCR product was subcloned into the Bam HI site in pUC19 vector by using HiFi DNA Assembly Cloning Kit (New England Biolabs), then pUC19-2+1-100-nt combiG4 model sequence construct was obtained.

**pUC19-2+1-140-nt combiG4 model sequence:**

The 2+1-140-nt DNA fragment was prepared by overlap extension PCR. Four dsDNA fragments were prepared by DNA polymerase reaction, using synthetic oligonucleotides [5′-CAG GTC GAC TCT AGA GTA ATA CGA CTC ACT ATA GAT CCT TTG AGG GTG GGT ATT CTC ATC TTA CCG CTG TTG AGA TCC AGT TCG ATG TAA CCC ACT CGT GCG CAG GAA AGA ACA TGT GAG CAA AAG GCC-3′] and [5′-GAG CTC GGT ACC CGG GTA ACG CGA ATC CCA GGT TCA GAA ACG CTG GTG AAA GTA AAA GAT GCT GAA GAT CAG TTG GGT GGG CCT TTT GCT GGC CTT TTG CTC ACA TGT TCT TTC CTG CGC-3′]. The dsDNA was subsequently amplified by PCR, using a forward primer [5′-CAG GTC GAC TCT AGA GTA ATA CGA CTC ACT ATA GAT CCT TTG AGG-3′] and a reverse primer [5′-GAG CTC GGT ACC CGG GTA ACG CGA ATC C-3′]. The PCR product was subcloned into the Bam HI site in pUC19 vector by using HiFi DNA Assembly Cloning Kit (New England Biolabs), then pUC19-2+1-140-nt combiG4 model sequence construct was obtained.

**pUC19-2+1-200-nt combiG4 model sequence:**

The 1+1-200-nt DNA fragment was prepared by overlap extension PCR. Four dsDNA fragments were prepared by DNA polymerase reaction, using synthetic oligonucleotides [5′- GGG TAT TCT CAT CTT ACC GCT GTT GAG ATC CAG TTC GAT GTA ACC CAC TCG TGC GCA GGA AAG AAC ATG TGA GTC GAC GCT CAA GTC AGA GGT GGC GAA ACC CGA CAG GAC TAT AAA GAT ACC AGG CG -3′] and [5′- AGG TTC AGA AAC GCT GGT GAA AGT AAA AGA TGC TGA AGA TCA GTT GGG TGG GCC TTT TGC TGG CCT TTT GGG AAA CGC CTG GTA TCT TTA TAG TCC TGT CGG G -3′]. The dsDNA was subsequently amplified by PCR, using a forward primer [5′-TAT AGA TCC TTT GAG GGT GGG TAT TCT CAT CTT ACC GCT GTT GAG-3′] and a reverse primer [5′-CGG GTA ACG CGA ATC CCA GGT TCA GAA ACG CTG GTG AAA GT-3′]. The dsDNA was subsequently amplified by PCR, using a forward primer [5′-CAG GTC GAC TCT AGA GTA ATA CGA CTC ACT ATA GAT CCT TTG AGG-3′] and a reverse primer [5′-GAG CTC GGT ACC CGG GTA ACG CGA ATC C-3′]. The PCR product was subcloned into the Bam HI site in pUC19 vector by using HiFi DNA Assembly Cloning Kit (New England Biolabs), then pUC19-2+1-200-nt combiG4 model sequence construct was obtained.

**Preparation of pIRES-combiG4 model seuqence-5′UTR-NL-FL Constructs**

1+1-100-nt or 2+1-100-nt combiG4 model-5′UTR DNA fragments were amplified by PCR using pUC19-1+1-100-nt combiG4 model sequence or pUC19-2+1-100-nt combiG4 model sequence constructs as templates by PCR-amplification with a primer set for 1+1-100-nt G4 combiG4 model [5′-ATA GGC TAG CCG ATC CTT TGA GGG TAT TCT CAT C-3′] and a reverse primer [5′-GAA GAC CAT GAA TTC TAA CGC GAA TCC CAG GTT CAG-3′], and a primer set for 1+1-100-nt G4 combiG4 model [5′-ATA GGC TAG CCG ATC CTT TGA GGG TGG GTA TTC TCA TC-3′] and a reverse primer [5′-GAA GAC CAT GAA TTC TAA CGC GAA TCC CAG GTT CAG-3′], respectively. Each PCR product was subcloned into the Xho I site in pIRES-NL-FL construct by using HiFi DNA Assembly Cloning Kit (New England Biolabs), and then pIRES-model-5′UTR-2+2-100nt-NL-FL constructs were obtained.

**Construction of Gs-Staple oligomer and As-Staple oligomer Expression Vectors**

**pAAV-mTRPC6-Gs-Staple oligomer or As-Staple oligomer:**

The dsDNA fragments of the 41-nt Gs-Staple oligomer and 41-nt As-Staple oligomer were prepared by annealing with synthetic oligonucleotides [5′-GAG AAA AGC CTC TAG ACA GGC GGC CCC AGG CTT GGG TGG GTT AGC AAA GCA ATG CCA TTT TTT TCT AGT GAT ATC GAT A-3′] and [5′-TAT CGA TAT CAC TAG AAA AAA ATG GCA TTG CTT TGC TAA CCC ACC CAA GCC TGG GGC CGC CTG TCT AGA GGC TTT TCT C-3′], and [5′-GAG AAA AGC CTC TAG ACA GGC GGC CCC AGG CTT AAA TAA ATT AGC AAA GCA ATG CCA TTT TTT TCT AGT GAT ATC GAT A-3′] and [5′-TAT CGA TAT CAC TAG AAA AAA ATG GCA TTG CTT TGC TAA TTT ATT TAA GCC TGG GGC CGC CTG TCT AGA GGC TTT TCT C-3′], respectively. Each dsDNA was subcloned into the Xba I-Spe I site in a pAAV-U6-ZsGreen1 vector (Takara Bio) using HiFi DNA Assembly Cloning Kit (New England Biolabs), and then pAAV-mTRPC6-Gs-Staple oligomer-41-nt and pAAV-mTRPC6-As-Staple oligomer-41-nt constructs were obtained.

**pAAV-hTRPC6-Gs-Staple oligomer:**

The dsDNA fragments of the 41-nt Gs-Staple oligomer and 41-nt As-Staple oligomer were prepared by annealing with synthetic oligonucleotides [5′-GAG AAA AGC CTC TAG ACA GGC GGC CCC AGG CTT GGG TGG GTT AGC AAA GCA ATG CCA TTT TTT TCT AGT GAT ATC GAT A-3′] and [5′- TAT CGA TAT CAC TAG AAA AAA CCA CTC GGC TCC CCC GGG AGA ACC CCA CCC CAA CTC CTC GGC CTG ACC TGC TCC CTA GAG GCT TTT CTC-3′]. The dsDNA was subcloned into the Xba I-Spe I site in a pAAV-U6-ZsGreen1 vector (Takara Bio) using HiFi DNA Assembly Cloning Kit (New England Biolabs), and then pAAV-hTRPC6-Gs-Staple oligomer-53-nt was obtained.

**Preparation of RNA templates for NMM fluorescence assay and thermal denaturation profiles**

The dsDNA for RNA transcription was prepared from pUC19-combiG4 model sequence, pUC19-mTRPC6-5′UTR constructs by PCR-amplification with a primer set for combiG4 model sequence [5′-CAG GTC GAC TCT AGA GTA ATA CGA CTC ACT ATA GAT CCT TTG AGG-3′] and [5′-CGG GTA ACG CGA ATC CCA GGT TCA GAA ACG CTG GTG AAA GT-3′], and a primer set for mTRPC6 5′UTR [5′-TAG AGT ACT TAA TAC GAC TCA CTA TAG GG-3′] and [5′-GGC ACA GTG CCT GGC CGG-3′], respectively. The dsDNAs were transcribed into single stranded RNAs (ssRNAs) using ScriptMAX® Thermo T7 Transcription Kit (Toyobo). The ssRNAs were purified with After Tri Reagent RNA Clean Up Kit (Favorgen).

**Preparation of RNA templates for RTase stop assay**

The dsDNA for RNA transcription was prepared from pUC19-combiG4 model sequence, pUC19-mTRPC6-5′UTR constructs by PCR-amplification with a primer set for [5′-CAG GTC GAC TCT AGA GTA ATA CGA CTC ACT ATA GAT CCT TTG AGG-3′] and [5′-CGC CAG GGT TTT CCC AGT CAC GAC-3′], respectively. The dsDNAs were transcribed into single stranded RNAs (ssRNAs) using ScriptMAX® Thermo T7 Transcription Kit (Toyobo). The ssRNAs were purified with After Tri Reagent RNA Clean Up Kit (Favorgen).

**Preparation of RNA Templates for In Vitro Translation**

The dsDNA for RNA transcription was prepared from pIRES-1+1-100-nt combiG4 model sequence-NL-FL, pIRES-2+1-100-nt combiG4 model sequence-NL-FL constructs by PCR-amplification with a primer set for [5′-CAG GTC GAC TCT AGA GTA ATA CGA CTC ACT ATA GAT CCT TTG AGG-3′] and [5′-TTA CGC CAG AAT GCG TTC GC-3′], respectively. The dsDNAs were transcribed into single stranded RNAs (ssRNAs) using ScriptMAX® Thermo T7 Transcription Kit (Toyobo). The ssRNAs were purified with After Tri Reagent RNA Clean Up Kit (Favorgen).
